# How biological cells including platelets and megakaryocytes decide complex problems fast but risky

**DOI:** 10.1101/2021.09.10.459594

**Authors:** Juan Prada, Johannes Balkenhol, Martin Kaltdorf, Özge Osmanoglu, Martin Stoerkle, Katrin Heinze, Georgi Manukjan, Harald Schulze, Thomas Dandekar

**Author notes:** equal contributing first authors (Juan Prada, Johannes Balkenhol, Martin Kaltdorf, Özge Osmanoglu). **Abbreviations:** NP, non-deterministic complete; P, always polynomial time taking; SAT, boolean satisfiability problem; MK, Megakaryocytes; HSC, hematopoietic stem cells; CDC2, cyclin dependent kinase 1; TP53, tumor protein p53; SRC, proto-oncogene, non-receptor tyrosine kinase; MYC, MYC proto-oncogene; JUN, Jun proto-oncogene; MYC, Myc proto-oncogene protein; ERK, Mitogen-activated protein kinases; MTOR, Serine/threonine-protein kinase mTOR; AKT, RAC-alpha serine/threonine-protein kinases; ITPR, Inositol 1,4,5-trisphosphate receptor; CXCR, C-X-C chemokine receptor; DVL, Segment polarity protein dishevelled homolog DVL; RUNX2, Runt-related transcription factor 2; NFkB, Nuclear factor-κB; MMP9, matrix metallopeptidase 9; Pi3K, phosphoinositide 3-kinase; SRC, proto-oncogene tyrosine-protein kinase Src; LYN, tyrosine-protein kinase Lyn; AKT1, RAC-alpha serine/threonine-protein kinase; TNFRSF1A, tumor necrosis factor receptor superfamily member 1A; TRADD, tumor necrosis factor receptor type 1-associated DEATH domain protein; ILK, Integrin-linked protein kinase; ITGB1, Integrin beta-1; GRB10, growth factor receptor-bound protein 10; CAC, cytosolic calcium release; G6b-B, megakaryocyte and platelet inhibitory receptor G6b; VC, value control centrality; DC, dynamic control centrality; TC, total control centrality; CDKN1A, Cyclin-dependent kinase inhibitor 1; CHUK, inhibitor of nuclear factor kappa-B kinase subunit alpha.

## Abstract

Mathematical decision processes are accurate but sometimes take very long time or simply do not happen. Decisions in biology happen fast and driven by evolution, optimizing survival chances. This results in stochastic decisions with on average good adaptation to the environment but an inherent risk of individual errors e.g. developing cancer during cell regeneration. We calculate and show in platelets and megakaryocytes how cellular decision processes increases risk for errors and inflammation. Short cut solutions adapted from nature improve computer strategies for protein folding and network decision processes. Complex problems are not always solved in foreseeable time, instead the fast solutions in biology speed up errors everywhere including biochemical aging of blood vessels, misfolded proteins, mis-programmed cells, cancer and heart failure.

**One sentence abstract:** We investigate in biological networks how complex decision problems are mastered not by an accurate but unforeseeable long mathematical search but rather pragmatic and fast, with an inherent risk of error, a basis for inflammation and cancer.

## Introduction

Complex decision processes are fundamental problems for computation. Well known are the non-deterministic complete or **NP**-problems [1] and one of the big challenges in mathematics is about whether there may be a mapping from these **NP**-problems to always polynomial time taking (**P**) problems, which are more certain and easier to solve. As a solution is highly contested but the answer to this is completely open [2], we shed light on complex decision processes from a biological perspective and look at the biological consequences. We show how in biological systems challenging decision processes (**NP** problems; [1] are often pragmatically treated using stochastic decisions, which on average optimize survival for the population and avoid stalling of decision processes. The latter can happen in formal systems, if fundamental statements have to be decided (Gödel’s incompleteness theorems, reviewed in [3]. We show that such fundamental decision problems occur in cell cycle progression, and that the stochastic resolution implies errors in cell division. Our finding could be an important factor for the increasing risk of cancer with every cell division (Tomasetti et al., 2017). Analyzing a similar network in hematopoietic stem cells (HSC) we show in detail involved network decisions and different types of decision processes involved in cancer and apoptosis. We next show how unsure decisions and unstable thresholds in platelets lead to a fragile balance of thrombosis and hemostasis but also imply thrombo-inflammation and myelofibrosis. Our analysis pinpoints new potential target proteins in platelet inflammatory processes (Stoll and Nieswandt, 2019).

**NP** complete calculations are in molecular biology nearly always reduced to simpler calculations in polynomial time. This allows us to suggest new heuristics to simplify an **NP** complete problem such as protein folding and to derive fast heuristic solutions, complementing current recent advances from artificial intelligence-based approaches of protein fold and energy potential recognition [5]: We adapt an efficient Maxcut algorithm [6] for constrained-based protein structure prediction e.g. from NMR data. However, in biological systems protein folding uses not a perfect search strategy, implying a risk for misfolded proteins. Mistakes in folding may lead with time to protein precipitation and Alzheimeŕs risk [7]. In general, molecular biology heuristics avoids **NP** calculations in order to allow fast decisions. This enhances the risk for biochemical accidents and cellular clock errors, the two major factors for aging [8]. Only evolution may instead tackle **NP** problems head-on using millions of years and large populations.

These considerations motivate our suggestion that in fact **NP** problems [2] are never reducible to P problems, relaxation always implies errors, and neglects (sometimes non-obvious) elements of complexity. Rejection of infinity and **NP** problems may seem mathematically possible (constructivism; Troelstra, 1977) but the true complexity of **NP** problems versus finite decision time manifests itself constantly in us by biochemical accidents and inevitable aging. In summary, the heuristics in biology cope well with otherwise unsolvable complex problems and inspire improved calculations, but the inherent errors imply a necessary risky decision in fundamental cellular processes, be it cell division, aging and cancer, platelet activation, resulting stroke and cardiac insult (Stoll and Nieswandt, 2019) as well as neurodegeneration, protein misfolding and Alzheimeŕs disease. As we perform for the first time large-scale simulations on platelet and hematopoietic cellular networks regarding such complex decisions processes, we pinpoint involved protein interactions, validate consequences by genetic, gene and protein expression data and expose key proteins involved for better pharmacological modulation and prevention of such disease states.

### Undecidable statements and their stochastic resolution in cell cycle

According to Gödel’s theorems, there are clear limits for decisions as well as completeness of formal descriptions of decision processes. The proof can be done by using self-referring statements (Gödel numbers representing not only numbers but also logical statements) and proving that the truth value cannot always be found. To mathematically mend such situations always one statement more is required *ad infinitum* [10].

We show here by direct translation of the statements of the proof that cellular decision processes can also have such self-referring statements (Box 1) since central genes (e.g. cyclin dependent kinase 1 (CDC2), tumor protein p53 (TP53), SRC proto-oncogene, non-receptor tyrosine kinase (SRC)) and resulting proteins represent not only the active enzyme part of the decision process but in such fundamental cell fate decision processes also a coding part of the process: they thus decide on the whole cell including the gene or protein itself. Such fundamental cellular decision processes are cell death (apoptosis), cell division etc. Here, no perfect decision is possible but conflicting statements pose a risk of a halt in the cellular machinery.

In <u>formal systems</u>, the critical step concerning the Gödel proof (see box 1 for explanation) is a statement such as:

> “Sentence G: This statement is not a theorem of TNT (Typographical Number Theory)”.

<u>In cellular systems</u> this translates into a statement such as:

> “This cell cycle state cannot be decided by the internal stored genetic model.”

**Box 1** shows how such statements look like if they occur in the cell. **Fig. S1** gives the underlying decision network for cell cycle and cancer related proteins. The involved complexity allows no shorter program [11]. Unlike mathematics, the decision process neither is in a loop or halts but is stochastically resolved, optimizing average survival probabilities but leading sometimes to serious errors. These errors increase linearly with the number of cell divisions, in line and supporting the observed linear increase of cancer risk parallel to the number of cell divisions (Tomasetti et al., 2017).

### Decision networks in hematopoietic stem cells and platelets imply cancer and thrombo-inflammation risk

We next studied the effect of decision errors in large networks. For the first time, we simulate here all decision processes concerning the network in large-scale models (see methods for network construction) of HSC (**Fig. S1**; 242 nodes and 519 interactions) and platelet (**Fig. S2**; 455 nodes and 821 interactions). We also show involved central protein network modules such as apoptosis, cell cycle, proliferation in HSC model and regulation of cell shape, and irreversible aggregation in platelet model (**Fig. 1a**&**b**, **Fig. 2a**).

**Figure 1:**
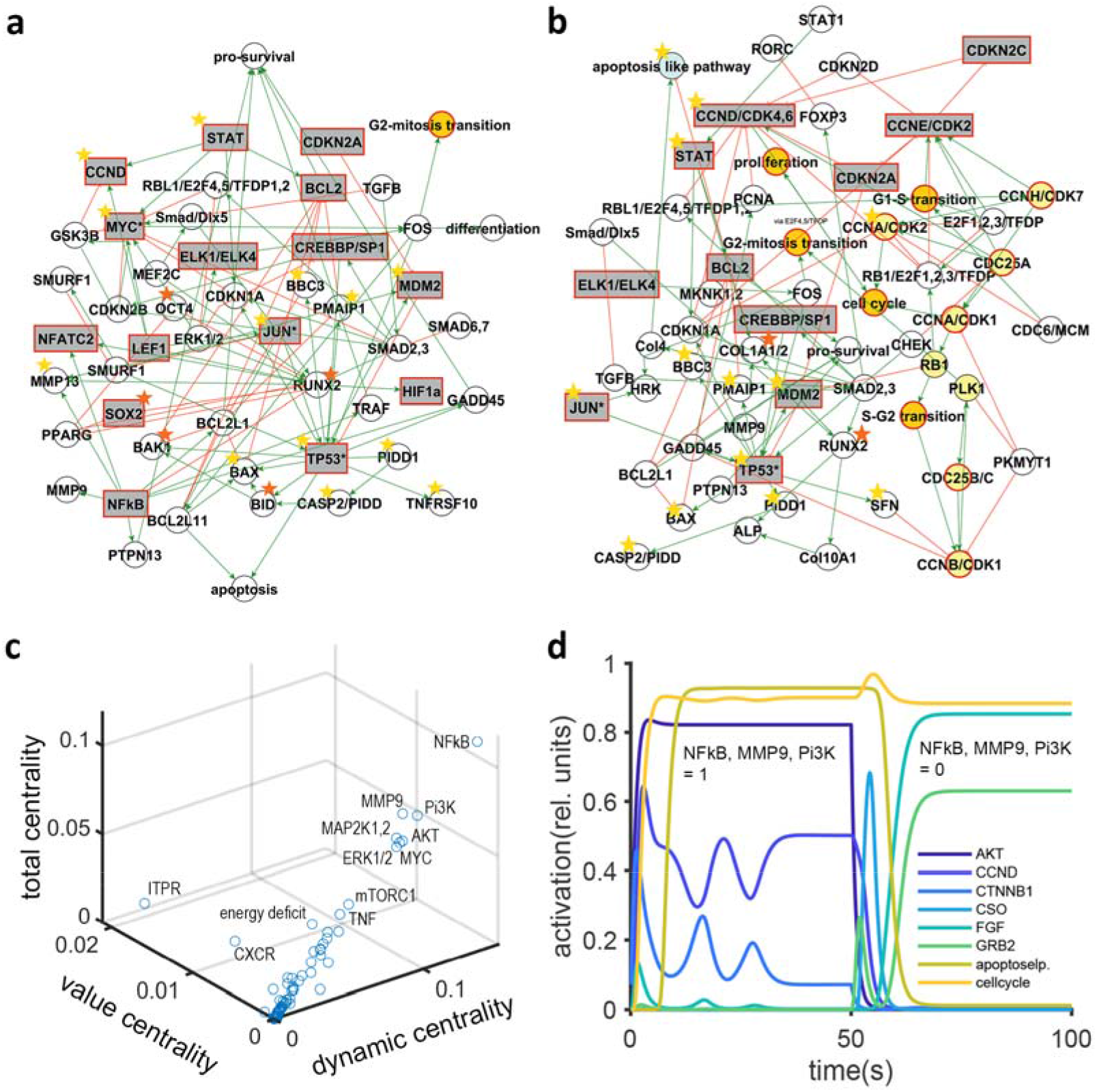
Network decisions and oncogenic processes in hematopoetic stem cells (HSC). **(a-b) In the HSC sub networks** (full network in Fig. S1) decision processes are influenced by cell cycle determinants and components (yellow fill) and cell cycle check-points (orange fill), HSC differentiation and suitable markers (pink), platelet interaction and differentiation factors (light blue), as well as general components of different signaling cascades and proteins for proliferation decisions mediating known oncogenic properties (grey colored rectangles). Undecided proteins are highly variable and change frequently the activation level when switching between stable states. The variable proteins are indicators for certain system states and are effectors of certain signaling pathways. We distinguish between those nodes undergoing clear decisions and unsure decisions. The stars represent clearly undecided (orange) or unsure undecided nodes (gold). **(a) Subnetwork on apoptotic processes** triggering platelet formation with error-prone decisions (red rims). **(b) Subnetwork on cell cycle** components and oncogenes. **(c) Network control in HSC**: direct, pharmacological control (value control centrality (VC; high: ITPR, CXCR, DVL)) is compared with dynamic control centrality (DC) of a protein via the network (high: NFkB, MMP9, PI3K,) and total control centrality (TC; high: NFkB, PI3K, MMP9). **(d) Influence of the high control proteins, such as NFkB, MMP9 and Pi3K on the HSC unsure decisions**. Activation of the central proteins to the maximum activation induces apoptosis and cell cycle changes (0-50 s). Subsequent deactivation of the same nodes deactivates apoptosis, but showed only minor influence on cell cycle (50-100 s).

**Figure 2:**
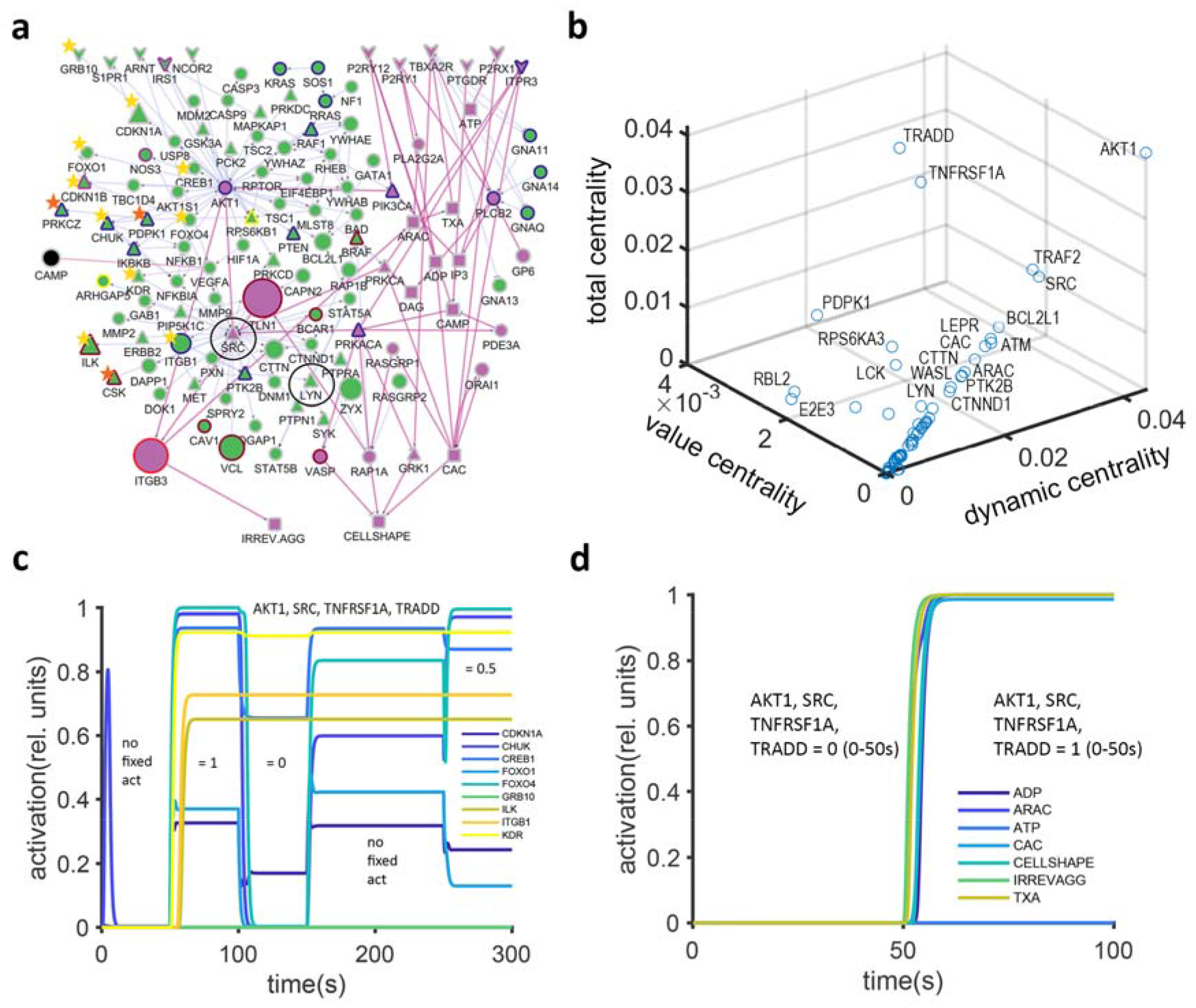
Unsure and clear decision processes in platelets. **(a) Platelet decisions proteins.** The stars represent clearly undecided (orange) or unsure undecided nodes (gold). The purple fill indicates the central regulatory cascade. The green fill of the nodes represents the first neighbors. Further proteins: kinases, phosphatases (triangles), receptors (arrow head), macroscopic effects, non-protein coding molecules (rectangle) and protein-coding molecules (circle) are underlined. Pathways are indicated in border colors (red: blood coagulation; blue: inflammation mediated by chemokine and cytokine signaling; dark red: integrin signaling pathway; pink: interferon-gamma signaling, interleukin signaling and plasminogen activating cascade; yellow: PDGF signaling pathway). The node size represents expression in absolute values of RPKM. **(b) Network control in the platelet** is indicated as in Fig. 1c. **(c) Influence of central proteins in platelets on unsure undecided nodes**. The top four DC proteins are tested on their influence on the unsure undecided nodes in (a). The activation levels of the top DC proteins are altered over time and the dynamics change of the unsure proteins activities are displayed. It is shown that the levels of, e.g. cyclin-dependent kinase inhibitor 1 (CDKN1A), inhibitor of nuclear factor kappa-B kinase subunit alpha (CHUK), ILK, ITGB1, are fine-tuned by the central proteins. **(d) Influence of high DC proteins on clear decisions.**

We analyzed the stability of the involved nodes along with the stability of the entire system. Our analysis revealed that the unstable decision nodes in HSCs lead either to successful maturation or sometimes cancer (major oncogenes MYC proto-oncogene (MYC), TP53 and Jun proto-oncogene (JUN); see excel file “oncogenes” for all considered here). These nodes are involved in the HSC protein network modules “cancer cycle” (**Fig. 1a**) and “cell cycle” (**Fig. 1b**). To reduce errors, nodes with high network centrality (**Table S1**) control decision nodes (**Table S3**) by integrating many pathways. We differentiate between decision nodes of high variance in activation level that change between two states (binary switch, or clear decision making) and decision nodes of high variance that change between many states (unsure deciders). Thus, a number of decision nodes have to be active and combined in the correct way for downstream transfer of information enabling diverse decision making.

For a more detailed analysis, key decision switches (i.e. network nodes that have high centrality) are compared (**Fig. 1c**, **Table S1**): This involves well known proteins (Myc proto-oncogene protein (MYC), Mitogen-activated protein kinases (ERK), Serine/threonine-protein kinase mTOR (MTOR), RAC-alpha serine/threonine-protein kinases (AKT)) that control and influence many other network nodes (DC, dynamic control) and proteins such as Inositol 1,4,5-trisphosphate receptor (ITPR), C-X-C chemokine receptor (CXCR), Segment polarity protein dishevelled homolog DVL (DVL) and Runt-related transcription factor 2 (RUNX2) that have high impact by direct control of their own activation (VC, value control) (Karl and Dandekar, 2015). Finally, in **Fig. 1d** we tested the influence of certain central nodes on unsure decision processes. Nuclear factor-κB (NFkB), matrix metallopeptidase 9 (MMP9) and phosphoinositide 3-kinase (Pi3K) are particularly labile tipping points changing system states, altering proliferation and apoptosis. Their activation induces apoptosis and cell cycle changes, whereas their deactivation inhibits these processes. The activation levels of these proteins are thus important since possible errors in cell cycle regulation increase directly (Tomasetti et al., 2017) cancer risk with every cell division.

On the other hand, central decision nodes in the platelet networks (**Fig. 2a**) such as proto-oncogene tyrosine-protein kinase Src (SRC) and tyrosine-protein kinase Lyn (LYN) and RAC-alpha serine/threonine-protein kinase (AKT1) are found to be well connected combining direct and dynamic network control (**Fig. S3**). Our large-scale calculations show different types of platelet decision nodes and their function within the network (**Fig. 2a**). As in the HSC network, we differentiate between clear decision making and unsure deciders. For a detailed analysis of the involved decision processes, **Table S3** (the stability analysis of the platelet network) presents and analyzes platelet decision modules *in silico*. This different types of nodes, decision behavior and network control are responsible for the fragile balance between platelet activation and inactivity, a fine-tuned physical response for thrombosis and hemostasis. However, there are different types of decision nodes (**Fig. 2a**), including mediators of thrombo-inflammation, signaling and blood coagulation.

For testing the influence of particularly unsure decision making, we annotated their pathway relevance. Our analysis shows (**Table S3E**) that on the cellular level unsure decisions involve platelet signaling, cytoskeletal interactions, cell division and apoptosis, while on the tissue level, this may stimulate chronic inflammatory processes such as myelofibrosis.

Similar to the analysis of the HSC network, the relationship of central proteins and hence decision nodes were investigated in the platelet network. The central nodes (**Fig 2b&S3**) such as SRC, AKT1, tumor necrosis factor receptor superfamily member 1A (TNFRSF1A) and tumor necrosis factor receptor type 1-associated DEATH domain protein (TRADD) can switch between stable network states. This gets clear when following up the activation level of unsure nodes in face of different activation levels of the central nodes (**Fig. 2c**). The above outlined integrin-linked protein kinase (ILK) and integrin beta-1 (ITGB1) show the downstream control by central nodes and thus are indicators for decision making. However, unsure decision making depends clearly on previous activations. This is indicated by ILK, or growth factor receptor-bound protein 10 (GRB10), that show different activation levels when the central nodes have the same state (0-50 s and 150-250 s). This indicates that unsure decision-making is context depended, complex and rather statistical.

Furthermore, platelets induce also clear decisions such as irreversible aggregation, cytosolic calcium release (CAC) and cell shape changes (**Fig. 2d**). The same central nodes that influence unsure decisions (**Fig. S3**), control clear decisions. The clear decisions rather rely on positive feedback, that is activated once a threshold is overstepped. There is a decreased likelihood of complex influence on the decision nodes beyond the threshold, thus some processes appear irreversible.

In order to follow up and validate the simulated decision processes in experimental data, **Table S3E** adds RNAseq data of megakaryocytes (MK) in a megakaryocyte and platelet inhibitory receptor G6b (G6b-B) knockout mice, leading to chronic myelofibrosis in mice. The observed phenotype and gene expression values compared variance in simulated activity, and the respective variance in expression in MKs. Certain unsure decision nodes (ILK, ITGB1) simulation showed differential regulation in the G6b-B knock-out study, especially when investigating male and female differences and also showed high variance in expression across all samples.

The unsure decision nodes such as ILK and ITGB1 appear in networks downstream of central control proteins balancing activating and inhibitory pathways (e.g. **Fig. 1a, b and 2a**), thus indicating heuristic decisions in face of complex integration. In comparison, the clear decision nodes, such as SRC, irreversible aggregation and cell-shape alterations (**Fig. 2d**) rely on clear threshold dependent dynamics, that translates complexity into a clear binary code.

The complexity is here reduced into the approximation of stable energy states of a network that encode e.g. the platelet response to stimulus (**Fig, 2a** and **Table S3**). The more complex a network gets, by adding more nodes, edges and feedback loops, the more stable states can be determined and thus the more complex responses on stimuli are possible. Unsure and undecided nodes, with their high variation and their many stable activation levels, are indicators for complex networks. Yet, with growing complexity it is more difficult to establish clear decisions. Overall, these are highly accurate decisions by the platelet network but still suboptimal, simply because it is not possible to better decide in such a fast time about an **NP**-complete problem.

### Molecular processes: Protein folding, aggregation and neurodegeneration

In molecular biology, **NP** complete calculations are often reduced to simpler calculations in polynomial time to achieve fast decisions. Protein folding, where proteins transform from primary structure into their tertiary structure is an often insurmountable challenge for **NP** calculations [13]. A whole repertoire of forces is involved such as hydrogen bonds, van der Waals interactions, electrostatic interactions [14], as well as hydrophobic interactions.

Fast biological decision processes happen efficient but heuristically in protein folding as well as in signaling as shown above. This suggests a new, heuristic solution strategy for identifying the optimal protein fold (**Fig. 3a**, code example in supplemental material) by treating this task as a graph Maxcut problem [15]: Find the partition of a graph such that two subgraphs are formed while maximizing the value of edges cut. Each amino acid in the protein sequence is modeled as a vertex in the graph and each weighted edge represents the estimated energy reduction if the forming vertex pair would end up bounded in the tertiary structure. This Maxcut search is **NP**-hard. Hence, a semidefinite relaxation is introduced as proof of concept and resolved using the Goemans-Williams algorithm with a lower bound accuracy of 83% for its interaction predictions, in this case residue-residue contacts. Moreover, this can be further improved by cautiously setting the initial conditions. In the case of the 2D lattice folding problem, we formulated a convex optimization solution, where basic restrictions are incorporated and the key factor to achieve the optimal solution is to properly set the initial conditions. The example code for this approach is presented in the supplemental material together with the 2D lattice results for a small globular protein. It can easily be transferred to optimize constrained-based 3D predictions on protein structures, e.g. from NMR and there still the lower bound for contact prediction holds.

**Figure 3.**
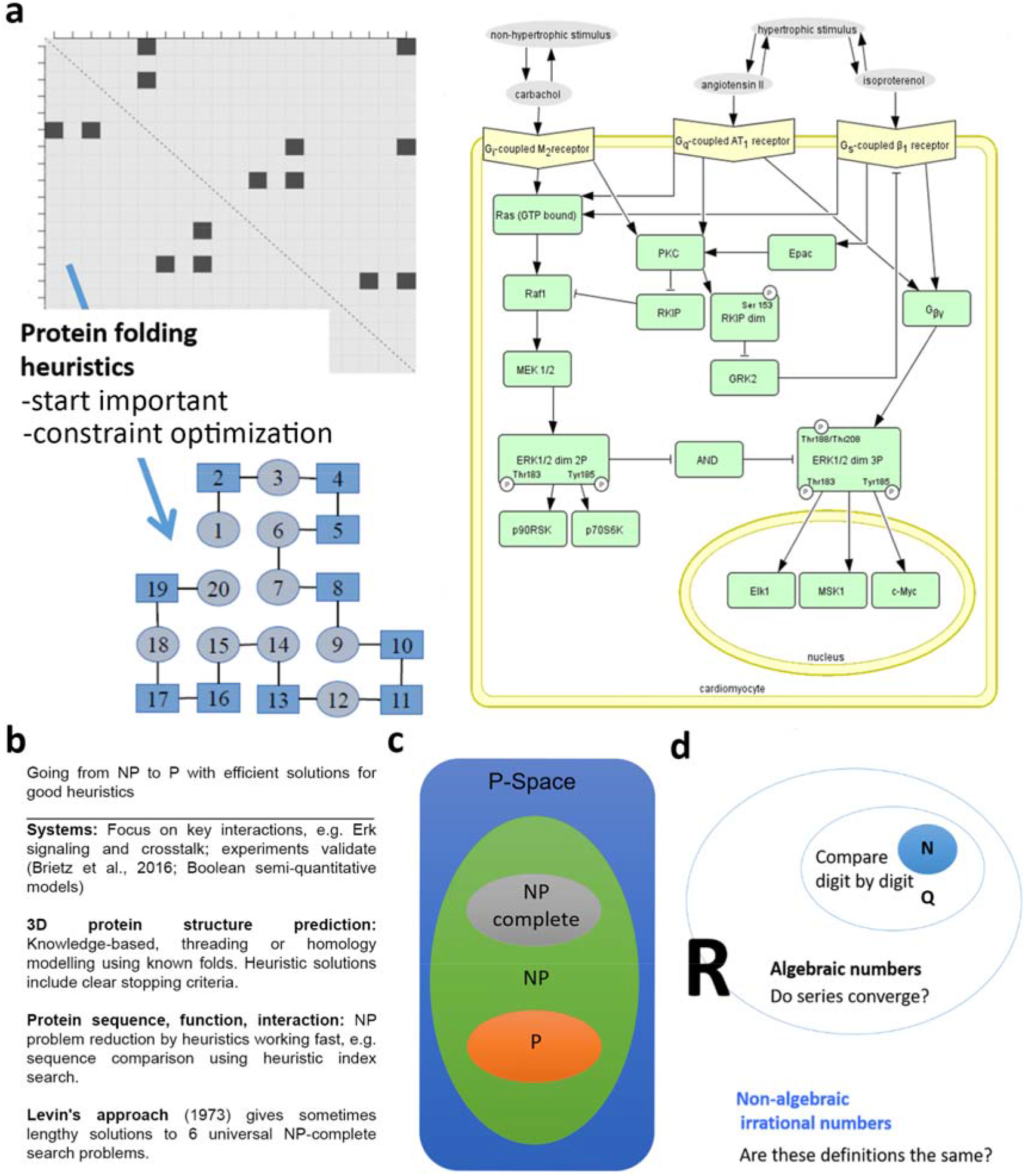
Biological heuristics may simplify some NP problems. **(a) Protein Folding approximately solved as a Maxcut problem. (top)** The primary structure of the protein can be presented as a full graph with n amino acids. Each is a vertex and each weighted edge encodes the free energy reduction during folding. (**bottom**) the Maxcut solution corresponds to find the set of edges that account for the biggest drop in free energy, a recursive **NP**-complete continuous model that cannot be efficiently solved. However, using the algorithm by Goemans and Williamson, (1995) the Maxcut problem can be solved with at least a 0.878 approximation. Nature seems to use such heuristics while folding proteins[16, 17]. **(b) The SAT equivalence holds also for signaling networks.** An example network is given (cardiomyocyte signaling, receptors on top). Decision processes can be modified to prevent cardiac hypertrophy. The proof for SAT-equivalence is given in supplemental material. **(c) 4 examples of efficient solutions and good heuristics from biology. (d) Comparing P and NP problems.** Shown are the relations of P and **NP** problems in problem space (P-Space, blue) according to [41]. In biology decisions have usually to be achieved fast and **NP** problems are reduced to **P** via heuristics. **(e) There may be no mapping possible from P problems to NP problems.** We suggest that some unpredictable long calculations stem from the fact that in contrast to natural or rational numbers as well as algebraic calculable numbers such as pi and e, many real numbers are not easy to distinguish from each other (the transcendent numbers).

Protein folding is rapid but not perfect in nature, instead of solving the **NP** problem it uses a shortcut, optimizing interactions (**Fig. 3a**, bottom) and by this energy and compactness, following folding funnels (Chen et al., 2019). Successful protein folding close to the lowest energy conformation depends on the start conformation (“start dependence”). An important aspect that adds to the complexity are chaperons helping protein folding [18] to prevent aggregates or folding start from an unfavorable conformation, since misfolded proteins may catalyze even more misfolded protein aggregates and lead to neurodegenerative diseases (see also **Table S2** in supplement).

Our algorithm is not tracing the complex cellular environmental conditions but stresses that initial conditions are important while later folding mechanisms relax the **NP** folding problem to a P problem applying also in nature heuristics to easily reach a close to minimum energetic conformation.

### Molecular biology heuristics often simplify complex decision problems leading to accumulation of errors and biochemical aging

All complex decision problems that are only non-deterministically solvable in polynomial time are equivalent (satisfiability (SAT) equivalence; Cook, 2009). This holds also for signaling networks (**Fig. 3b**). This was never shown before. We prove (see supplement) that the decision process in a biological signaling network (like stem cell or platelet networks) is analog to the well-known SAT problem in computer science. However, each biological module is in addition not mathematically accurate but heuristic, has only few nodes and this ensures that there is fast decision in polynomial time. The robustness in the platelet signaling applies only to the different signaling modules, these are failure tolerant and redundant and robust, only if central hubs are affected, their function stops (**Table S1** in Supplement).

In cellular decision processes, there seems to be never anything coming close to an **NP** problem. In particular, there is no unforeseen decision delay but fast heuristics in which combination of values in a Boolean function will result in a true or false value. In signaling networks something similar is happening: the cell has to decide which combination of active/inactive proteins will result in the desired response. This has to be decided fast and in most of the cases it involves a big network, in which reaching an optimal solution is challenging. The organism is trained to make accurate guesses on each situation which most of the time turn out to be correct but with no guarantee of optimality at all. However, appreciating this shows again that the cell has to cope with really tough problems and as they need to be stochastically resolved, this implies again an unavoidable risk of errors in our cellular decisions.

This motivates our suggestion that in fact **NP** problems are even mathematically never reducible to P problems, relaxation always implies errors. The complexity behind the non-deterministic behavior is not always obvious. However, there are good heuristic solutions for biological systems (**Fig. 3c**) that relax tough **NP** problems to simpler **P** problems (**Fig. 3d**), implying swift and well adapted cellular reactions as well as an inherent risk for errors. The latter includes biochemical accidents and impairs cellular clocks, major biological processes for aging.

#### P vs NP problems compared: number comparison example

Is it possible to use a computer to efficiently (polynomial time) solve problems which can otherwise only be solved efficiently by an omniscient oracle? We think that all problems that create a clear description of the output can always be searched and also printed out in polynomial time **P**. Hence, we think that all **NP**-hard problems have a non-obvious element of opacity in their definitions. It would be helpful to reveal this clearly: Once done, it then becomes clear that no mapping from the **NP** problem to **P** is possible. This conjecture is illustrated for the problem „denote two number representations for the same number?“ (**Fig. 3e)**. Regarding rational numbers this is a **P** problem: for instance, using the decimal representation just the digits of two numbers have to be compared to decide about similarity or not. However, for real numbers (and all more complex numbers), representations have to be compared which can be limits of different infinite series and there may be even several definitions potentially available such as different series converging and describing the transcendental number pi or, after all, something different from it. Finding the solution to whether two such representations describe the same number takes sometimes unexpectedly long (this is again probably an **NP** problem, as equivalent to a SAT problem such as the subset sum problem, see supplemental material).

#### Conjecture 1

A general mapping from the non-deterministic polynomial time requiring complex **NP** problems to the clearer, only polynomial time requiring **P** problems is in fact only artificially possible, if you ban higher complexity from the “allowed” problems using “constructivism”, a specific non-mainstream school of mathematics [9] which allows to discard ill-defined entities (see below). Similarly, we suggest only if we ban in general all non-trivial potential alternative descriptions (requiring non-deterministic time to find out) for the same solution, then the non-deterministic behavior of solution finding will stop. These would be similar route lengths in the classical travelling salesman problem. In our toy example “number comparison” this corresponds to rejection of actual infinity (“constructivism”) as numbers which differ only after an infinite number of digits are then banned. The mathematical school of constructivism implies to reject other things in addition as not correctly defined, for instance the indirect proof or the different cardinalities of infinity after the mathematician Cantor.

#### Conjecture 2

More carefully phrased, we suggest that the challenge of **NP** problems in general is that there are multiple representations for some of the solution elements possible and that for these multiple representations it is non-determined how fast you realize that these are the same solution. Accordingly, our **conjecture 2** is hence that due to this more general reason, you can believe that a clear mapping to **P** problems is *not* possible.

#### NP problems are real and have to be mastered in cellular life

Rejection of infinity and **NP** problems may seem axiomatically possible (constructivism, see below; Troelstra, 1977) but the true complexity of **NP** problems in our real world versus finite decision time for biological processes manifests itself constantly in us by biochemical accidents and inevitable aging. Errors enhance the two major factors involved in aging (Sergiev et al., 2015), cellular damage by wrong decisions and protein misfolding as well as mis-programmed cellular clocks and maturation processes. The latter was strikingly shown by reprogramming [20]. **NP** problems doubtless exist in nature and are tackled head-on in biology just by evolution with millions of individuals over long time in parallel or in numerous instances in physics.

Polls among mathematicians show a clear majority in favor of **P** being different from **NP** [2]. Humans “have an understanding of concept of infinity” [21]. Another way to phrase the capabilities of living observers is that they have a concept of meaning and can hence organize decision processes efficiently for survival though the environment is complex and potentially infinite. Hence, they use again evolutionally driven, efficient and somewhat error-prone biology and which a machine does not share and hence cannot cope with.

Cellular decision processes typically reduce **NP** problems to many separated **P** problems, as represented by the various pathways downstream of receptors (platelet and HSC pathways). As the pathways are now separately analyzed, the cell network needs to integrate the signals again to recreate the **NP** complexity, of the analyzed several **P** complex pathways. In the network we see integration point such as AKT1, SRC and cytosolic calcium (CAC) that integrate different downstream pathways and transmit the integrated information to decision nodes (**Fig 1d**, **Fig. 2d**). Individual decisions such as receptor signaling or functional responses are coupled by integrative nodes. On these upstream nodes, the complexity of the world is reconstructed. The decisions are tested on the world and coupling parameters adjusted on that subsequent feedback. The coupling of the nodes in the network, the decision of the analyzed principle components of the decision view and the reconstruction of the lost initial condition due to dimension reduction are all potential sources of errors in the process of heuristic analyzing the **NP** complex world.

#### Conclusion

In our everyday world, complex problems may take unexpectedly long to solve (**NP** problems), there is no simple relaxation to **P** problems possible. Biological systems have not sufficient time to dwell on such complexities, use heuristics and have instead to cope with a certain amount of errors. Analyzing cellular decision problems and protein folding we show here in detail how decision processes nevertheless happen fast and direct in protein folding and cell cycle, platelets, and hematopoietic stem cells. However, this implies a natural risk for error, resulting in protein aggregates and inflammation as well as major diseases and causes of death such as cancer, neurodegeneration, and stroke. As a concrete demonstration megakaryocyte and platelet network properties are calculated and tested in a megakaryocyte knockout leading to myelofibrosis.

## Supporting information

Supplemental Table 2 Centralities Platelet CC

Supplemental Table 3 Stable State Analysis

Supplemental Table 3E Undecided Nodes in RNAseq

Supplemental Table 5 Network Platelet CC

Oncogenes

Supplemental Table 1 Centralities HSC

Supplemental Table 4 Network HSC

## Author contributions

JPS performed the protein folding simulations and found and developed the Maxcut strategy. JPS provided expert advice on NP problems and computer science and showed the SAT equivalence for cellular decision problems. JB performed the large-scale platelet and HSC simulations and bioinformatically analyzed all transcriptome and proteome data including the G6Bb knockout in megakaryocytes. MK, KH and ÖO provided systems biological expertise. MS provided engineering expertise. GM and HS performed the megakaryocyte and transcriptome experiments from GP6b knockout and wild type. HS provided platelet and megakaryocyte expertise. JPS, JB and TD drafted the manuscript. JB and JPS were supervised by TD. TD conceptualized, led and guided the study. All authors (JPS, JB, MK, ÖO, MS, KH, GM, HS and TD) edited the manuscript, gave comments and agreed to its final version.

## Data availability

All data for this study are available from the manuscript, its figures and its supplements. Simulation details and data are provided in full. All omics data for validation have been deposited and access identifiers given, in particular the set GSE155735 (transcriptome of native murine megakaryocytes; wild type and G6b-B knockout).

## Acknowledgements

we gratefully acknowledge funding by DFG (Project number 374031971 – TRR 240/Z2; [for JB, KH: /INF; for HS: /A03]) and the Land Bavaria (core support).

## Online Methods

### Gödel decisions in cell cycle

The Box 1 followed the Gödel proof concept by Douglas Hofstadter [10] and translated it into cell cycle components using Cytoscape repositories of cell cycle (see supplement). The error prone cell cycle decisions shown in supplement imply a linear increased risk of cancer by cell division as observed (Tomasetti et al., 2017).

### Network analysis of decision processes in platelets and hematopoietic stem cells

We first established the involved networks according to known protein databases of protein-protein interactions (in particular the PlateletWeb [22], the central activating cascade of mouse and man [23] and further literature (see **Table S5**) and inference methods [24]. **Table S4** lists all proteins involved in the hematopoietic stem cells (HSC) network and related literature. **Table S5** lists the platelet interactions of the central cascade (CC; according to [23,25–29]. The network is extended to the 3^rd^ neighbor using PlateletWeb [22] and combined with an inference method [24]. In addition to PlateletWeb, the platelet proteins are verified by RNASeq datasets [23,24,30]. We used the Uniprot protein [31] notation, for symbols and full names in the supplementary tables, in the figures and in the text.

The visualization of the networks (https://tr240.uni-wuerzburg.de/vippclass/index.php/s/roP72PoXzZjyQKf) is done with Cytoscape [32] and the yEd graph editor (https://www.yworks.com/products/yed). The pathways were annotated according to the panther pathway code [33].

Next, the software Jimena (https://www.biozentrum.uni-wuerzburg.de/bioinfo/computing/jimena/) allowed to dynamically simulate each network and identify system states, fragile decision points and different types of network control for each decision node: direct control is exerted by high “value centrality” (VC) and dynamic control over neighboring network nodes, so called “dynamic centrality” (DC) as well as combined control, i.e. “total centrality” (TC). The method and mathematical definition are described in detail in (Karl and Dandekar, 2015).

Large-scale simulations of decision nodes and system behavior were run on a workstation (40 twin cores). The run time was up to 48 hours; there were 10.000 trials on each of the networks, i.e. HSC with 242 nodes and 519 interactions, platelets with up to 455 protein nodes and 821 interactions.

Node behavior was classified into the categories shown in the results figures: All three centrality measures as well as determinism of the decision: Clearly stable nodes (SRC), undecided stable nodes and undecided unstable nodes. We did run extensive simulations for system stability, stable and unstable decision nodes: The networks had sizes of 242 nodes (hematopoietic stem cell network) and 455 nodes (platelet network). For each network 200 simulations were sampled. The activation of unsure and clearly undecided nodes was tested in Jimena in response to changes of stable nodes with high dynamic centrality, such as AKT1 and SRC. For instance, SRC is a bi-stability switch and this bi-stability switch property is analyzed and experimentally validated in detail in Mischnik et al. (2014) and appropriately modeled in our simulation. ITGB1 and ILK are under this condition unstable.

**Table S1** and **S2** indicate the calculated centralities of the network nodes. The centralities are visualized with Matlab 2019 (https://www.mathworks.com/products/matlab.html).

**Tables S3E** lists specific differences in mRNA expression and compares them for strong decision nodes (high centrality values) and unsure decision nodes as well as network connectivity parameters compared. Further columns of **Table S3** list the stable system states for the platelet and the HSC network. The nodes are ranked according to their variance of the activation level. The activation level ranges between 0 and 1. The activation level were separated in 10 states between 0 and 1 with a step width of 0.1. It was counted how many different states a node adopts. The nodes that show a high variability of states between 0.1 and 0.9 are called unsure undecided nodes. Nodes that show a high variability but switch between values >0.9 and <0.1 are determined as clearly undecided nodes. The cutoff of variance for undecided nodes was 10^-4^.

### RNAseq analysis

For RNAseq, native murine megakaryocytes (MKs) were isolated from bone marrow of tibiae and femora by centrifugation. MKs were isolated using anti-CD61 microbeads according to the manufacturers’ protocol (Miltenyi Biotec). CD61-positive cells were subjected to a BSA density gradient separation, centrifuged at 300 g for 5 min and lysed in 500 µl Trizol reagent (Invitrogen). RNA extraction was performed using the RNeasy Mini Extraction Kit (Qiagen). RNA quality was proven by a 2100 Bioanalyzer with the RNA 6000 Pico kit (Agilent Technologies). DNA libraries were prepared from 50□ng of total RNA with oligo-dT capture beads for poly-A-mRNA enrichment using the TruSeq Stranded mRNA Library Preparation Kit (Illumina). Sequencing was performed on the NextSeq-500 platform (Illumina) in single-end mode with 75 nt read length. Sequencing data are available at NCBI GEO (http://www.ncbi.nlm.nih.gov/geo) under the accession number GSE155735.

For differential analysis the gene raw counts were normalized with the R package RUVIII [37]. Male and female genes were tested for differential expression in the G6b-B deficient model with edgeR version 3.32.1 [38]. The genes were identified as differentially expressed when they showed a significant (*p*-value <0.05) change in read counts after multiple testing correction [39]. The variance of the gene expression was analyzed over all samples in order to compare it to variable genes identified in the network simulation.

### Protein folding heuristics

The Maxcut routine for protein folding followed Goemans and Williamson (2004), which achieves with high reliability a close to optimal solution. This is applied to a constraint-based 2D folding optimization algorithm. The strategy is easily transferred to 3D and constraint-based optimization of structure predictions from NMR data. Following our insights on protein folding simulations we find start value dependence both for the Maxcut approach as well as for protein folding (see details in supplement). The latter may lead to protein misfolding and protein aggregation as observed experimentally, too [40]; Table S2 in supplement)

### Extended figure description

**Figure 3. Biological heuristics may simplify some NP problems. (a) Protein Folding prediction optimized as a Maxcut problem. Top:** The primary structure of the protein can be presented as a full graph with n amino acids. Each is a vertex and each weighted edge encodes the free energy reduction during folding. Example: R for arginine, D for aspartic acid and a hypothetical interaction of 50 cal/mol. The graph has *n*(*n* − 1)⁄2 edges. The Maxcut problem splits the graph in two sub-graphs, the splitting passes through the edge combination with the highest weight. **Bottom:** the Maxcut solution corresponds to find the set of edges that account for the biggest drop in free energy, a recursive **NP**-complete continuous model that cannot be efficiently solved. However, using the algorithm by Goemans and Williamson, (1995) the Maxcut problem can be solved with at least a 0.878 approximation, hence at least 87,8% of the contact constraints are accurately predicted applying this strategy to constraint-based modelling of protein structure for instance of NMR data. Nature seems to use such heuristics while folding proteins[16, 17]. The amino acid chain is not rigid and accepts changes while still folding into the same structure. This is a form of relaxation like in semi-definite programming [15] allowing such a solution strategy including random movements and vibrations as observed in protein folding.

**(b) The SAT equivalence holds also for signaling networks.** An example network is given (cardiomyocyte signaling, receptors on top). Decision processes can be modified to prevent cardiac hypertrophy (3^rd^ Erk phosphorylation). The proof for SAT-equivalence is given in supplemental material.

**(c) Efficient solutions and good heuristics from biology.** The table gives four examples.

**(d) Comparing P and NP problems**

Shown are the relations of P and **NP** problems in problem space (P-Space, blue) according to [41]. In particular, even a quantum computer does not allow to solve all **NP** problems, it efficiently solves factoring and discrete logarithm calculations, however, graph isomorphism and **NP** complete problems (top, green circle) are still a challenge. In biology decisions have usually to be achieved fast. This happens often in polynomial space (P, red circle) but for other problems in biological systems, the complexity of the decision processes is high (**NP** problems, orange). Biological heuristics allow faster computations. For implementation in algorithmic strategies quite often not the **NP** problem is tackled but a reduction to a P problem covers most solutions relevant in practice. The largest space is **P**-Space, all problems solvable by a conventional computer with polynomial amount of memory but exponential number of steps.

**(e) There may be no mapping possible from P problems to NP problems.** We suggest that some unpredictable long calculations stem from the fact that in contrast to natural or rational numbers as well as algebraic calculable numbers such as pi and e, many real numbers are not easy to distinguish from each other (the transcendent numbers). Only by these additional, infinite dense and difficult to distinguish numbers the set of R has higher cardinality then N or Q. However, this leads to an unforeseeable behavior regarding the time required to calculate these or even just to print them in a conclusive way, distinguishing them from each other. We claim that all problems that create a clear description of the output can always be searched and also printed out in polynomial time **P**. In contrast, we think that all **NP**-hard problems have a hidden element of opacity in their definitions. This may however be difficult to pinpoint apart from the illustration problem to compare two definitions for a real number R and predict in advance the time required to decide whether they are the same or not. We conjecture that this last problem may also be an **NP** problem, but for that the equivalence of this “same definition problem” e.g. to the subset sum problem (**NP** proof for this see [42]) has to be shown. Nevertheless, at least this illustrate that a hidden element of opacity easily creeps up in **NP** problems and this may always prevent a clear mapping of **NP** problems to P and prevent a reduction of this larger problem class. Our conjecture may stimulate mathematicians to look at other **NP**-hard problems and rethink involved basic concepts such as the continuum hypothesis so that they become clearer defined allowing to achieve better calculation strategies with a clearer halting behavior. More details are given in Box M (mathematical problems and decision processes) and appendix D (mathematical considerations).

## Box I. Gödel statements occur in biological decision processes without stalling

We briefly summarise the Gödel proof for decision limits in formal systems as given by [10] (see also [43]). Next, we illustrate that cellular systems have the same self-referring capabilities (self-reference including arithmoquination capabilities) but nevertheless can deal with such system states. Finally, biological implications of such decision processes are discussed.

**1. Express with simple symbols all types of logical statements.**

<u>Formal systems:</u> Here you can use for instance TNT (Theoria Numerorum Typographica) a shorthand to express mathematical statements using simple symbols ([10],[43]; see also https://github.com/jeremyhuiskamp/tnt for a library of manipulating strings according to TNT). There are standard mathematical symbols such as +(plus), * (multiply), and =(equals). There are *variables,* represented by the letter a followed by primes: a, a’, a’’, *etc.* There are standard logical symbols such as ∼ (not), V (or), E (there exists) and A (for all). Finally, there are numbers, which are represented by the two symbols 0 (meaning zero) and S (meaning “the successor of”); so we count 0, S0, SS0, SSS0, and so on.

Starting from a list of axioms such as the Peano axioms (see [10],[43])

Axiom 1: Aa:∼Sa=0
Axiom 2: Aa:(a+0)=a
Axiom 3: Aa:Aa’:(a+Sa’)=S(a+a’)
Axiom 4: Aa:(a*0)=0
Axiom 5: Aa:Aa’:(a*Sa’)=((a*a’)+a)

Any type of proof and sentence on natural numbers can be formulated, e.g. ∼Ea:a*a=SS0 would be translated as “There is no square root of two.” A sentence or statement is *true* if it can be formed from the axioms. If the statement is *false,* we can derive its converse from the axioms. A Gödel statement G has a truth value which cannot be decided from the system though it can be derived from the system. Now we translate this proof here in this paper for cellular systems:

<u>Cellular systems:</u> Any type of cellular process can be encoded by genes which in turn yield gene products. Operations such as plus and multiply happen already on the nucleotide level. Variables are in this context genes and gene names, for these the standard logical symbols such as NOT active, OR (one of two gene products), E (there exists a gene such as …) and A (for all genes applies …) also exist.

**2.** <u>Formal systems:</u> The **critical step** concerning the Gödel proof is a statement such as: “Sentence G: This statement is not a theorem of TNT.”

<u>Cellular systems</u>: Here this would translate into a statement such as: “This cell cycle state is not in the internal stored genetic model.”

**3.** <u>Formal systems:</u> For the Gödel proof a **Gödelization** (give Gödel numbers to statements) is necessary to achieve self-referring states[43].

| Mathematical Logic | TNT | Gödel-numbered TNT |
| --- | --- | --- |
| An <u>axiom</u> is an "obvious" statement about natural numbers. | An <u>axiom</u> is a statement string | An <u>axiom</u> is a number |
| A <u>rule of production</u> is a logical way to work with axioms. | A <u>rule of production</u> is an allowed string-manipulation mechanism. | A <u>rule of production</u> is an allowed mathematical function. |
| The <u>theorems</u> you produce are | The <u>theorems</u> you produce | The <u>theorems</u> you produce |
| new statements about natural numbers. | are new strings. | are new numbers. |

**4. Finally, Writing Sentence G:**

a) first incomplete effort

T: The arithmoquine of A is not a valid TNT theorem-number.

A: Sentence T is not a valid TNT theorem-number.

However, this sentence A is not the sentence G we’re looking for, since it isn’t about itself, it’s about sentence T. We thus put:

T: The arithmoquine of A is not a valid TNT theorem-number.

A: The arithmoquine of Sentence T is not a valid TNT theorem-number.

*b) Hence, better and more accurate:*

**G: The arithmoquine of “The arithmoquine of A is not a valid TNT theorem-number” is not a valid TNT theorem-number.**

<u>Cellular systems</u>: In cellular systems the internally stored information also starts from genetically encoded axioms (so certain genes or sets of genes set basic circuits and internal values). A rule of production is a certain type of genetic circuit, and the combination or interference of genetic circuits produces more complex genetic circuits. The last sentence makes it clear that the Gödelisation and also the arithmoquination happen on the level of genetic circuits, in particular genetic circuits also deal with the state of the complete cell as a whole and / or can refer to themselves while describing internally the state of the whole cell, too (e.g. in cell cycle, Fig. 1S), e.g. a cellular representation of

*G: The arithmoquine of “The arithmoquine of A is not a valid TNT theorem-number” is not a valid TNT theorem-number.* would be

> “The cell cycle state referring to cell cycle control of G_1_ to S by activity of cdc25 on cdc2 is blocking the cell cycle control inhibition regarding transition of G_1_ to S activated by wee 1 mediated cdc2/cyclin B kinase phosphorylation which is itself controlling activity of cell cycle state.”

## Supplementary Material

### 1. In a nutshell: Why NP problems are tough to simplify?

To start with, there is the equivalence of all **NP**-complete problems (Cook’s theorem; [1, 2]). The Boolean satisfiability problem (SAT) is **NP**-complete and any other problem in **NP-**complete can be reduced in polynomial time to the SAT problem, using a deterministic Turing machine. Hence, once you could solve the SAT problem in **P** you can solve all **NP**-complete problems in **P** applying the theorem of Cook.

#### Conjecture 1

A general mapping from the non-deterministic polynomial time requiring complex **NP** problems to the clearer, only polynomial time requiring **P** problems is in fact only artificially possible, if you ban higher complexity from the “allowed” problems (using “constructivism”, a specific non-mainstream school of mathematics, which allows to discard ill-defined entities, see below). Similarly, we suggest only if we ban in general all non-trivial potential alternative descriptions (requiring non-deterministic time to find out) for the same solution, then the non-deterministic behavior of solution finding will stop. These would be similar route lengths in the classical travelling salesman problem. In our toy example “number comparison” this corresponds to rejection of actual infinity (“constructivism”) as numbers which differ only after an infinite number of digits are then banned. The mathematical school of constructivism implies to reject other things in addition as not correctly defined, for instance the indirect proof or the different cardinalities of infinity after the mathematician Cantor.

#### Conjecture 2

More carefully phrased, we suggest that the challenge of **NP** problems in general is that there are multiple representations for some of the solution elements possible and that for these multiple representations it is non-determined how fast you realize that these are the same solution. Accordingly, our **conjecture 2** is hence that due to this more general reason, you can believe that a clear mapping to **P** problems is NOT possible.

#### Application point

Moreover, from an application point of view we see that in reality NP problems are so challenging that they are really tackled in biology only in evolution with millions of years and non-deterministic solution finding behavior, but for all fast processes instead heuristics prevail. These allow fast decisions but imply a low probability for error. These may sometimes lead to important disease such as cancer, stroke but also aging and misfolded proteins. Hence, we think that mathematical constructivism is possible, but that this is not the correct model to describe the challenges of decision processes in biology, there clearly the NP problems are tough and not correctly solved but rather error-prone heuristics used. Similarly, NP problems doubtless exist in Nature and are tackled head-on by evolution with millions of individuals over long time in parallel in biology as well as in physics.

**The other parts of our paper build on these insights and show biological consequences from this:**

2. Cell cycle errors result from Gödel problems occurring in the cell (self-referring statements in cell cycle). It has been suspected earlier that living beings can solve such problems, but we show here for the first time a literal translation of the Gödel incompleteness proof into cell cycle division statements. Furthermore, the genes involved are well known cell cycle genes and interactors. So this shows for the first time that the observed linear increase of cancer risk with the number of cell divisions (Tomasetti et al., 2017, Tomasetti and Vogelstein, 2017) is unavoidable for the cell, as living beings have to cope with such Gödel problems / fundamental decisions and can only stochastically resolve these.
3. For hematopoietic stem cells the cell cycle considerations are shown in their effects in large-scale simulations never attempted before (see methods section). We furthermore show that in cellular decision processes different types of node determinism and stability can be distinguished as well as different types of network control. In particular, in addition there is an element of additional fragility by undecided decision nodes that can tip into a certain direction, in particular this concerns well-known proteins such as STAT, CDK4, JUN, TP53 in agreement with a strong literature on their importance.
4. For platelets this implies besides risk from cell division (Tomasetti et al., 2017) for hematological cancer and risk from unstable decision nodes (see 3) that we have also an inherent risk for fragile balance suddenly tipping. This is most evident in this cell type as platelets are at the route of thrombo-inflammation which then can lead to stroke and cardiac insult. For the medical important platelets and the HSC generating them we calculate for the first time the large-scale decision networks involved. Our simulations in the paper show the actual behavior of stable and unstable decision nodes. The unstable decision nodes are at the root of thrombo-inflammation caused by platelets and myelofibrosis in megakaryocytes. Moreover, these results are validated in detail by phenotypes from knockouts in mice gene expression data analyzed here. The fast decisions by the platelet protein network are still suboptimal, simply because it is not possible to decide in such a fast time an NP-complete problem.
5. Start dependence of protein folding. This is the consequence of biology applying only heuristics for finding the optimal protein fold. The start dependence implies that with a wrong start the protein folds incorrectly. In particular, incorrect folding and protein aggregates are possible, leading to neurodegeneration. Moreover, these considerations allow us to introduce a maxcut algorithm problem solving strategy for protein fold prediction. We show in 2D models that this strategy works and leads to good solution quality. By applying the Goemans-Williams algorithm for max-cut problem we can be sure to achieve an accuracy of at least 83% regarding correct interconnections, in this case residue-residue contacts for the protein. Moreover, this can be further improved by cautiously setting the initial conditions. We observed that nature folds protein extremely fast because it has learned to solve the problem as a convex problem, which actually keeps in a small vicinity within the funnel land scape. Nature carefully choses the initial conditions that lead to the desired configuration of the folded protein, we pretend to imitate this selection of initial conditions and observed that the problems becomes much simpler that thought before (from NP-complete to NP). We show this only for 2D toy examples which are easier to treat, but the limits and considerations set by the maxcut approach apply for all problems tackled including protein folding in three dimensions. So, the reported accuracy is a proof of concept on protein structure prediction which still has to be transferred to 3D predictions, for instance applying distance constraints to underdetermined 3D structure predictions from NMR data.
6. SAT equivalence of cellular decision problems. This was also never shown before. What we proof here is that, the decision process in a biological signaling network (like stem cell or platelet networks) is analog to the well-known SAT problem in computer science. The SAT problem consist in guessing which combination of values in a Boolean function will result in a true or false value. In the signaling networks a similar thing is happening, the cell has to decide which combination of active/inactive proteins which result in the desired response. This has to be decided fast and in most of the cases it involves a big network so reaching an optimal solution is not possible. The organism is trained to make accurate guesses on each situation which most of the time result correct but with no guarantee of optimality at all. However, appreciating this shows again that the cell has to cope with really tough problems and as they need to be stochastically resolved, this implies again an unavoidable risk of errors in our cellular decisions.

### 2. Full Hematopoietic Stem Cell network with subnetworks shown in Figure 1

**Figure S1:**
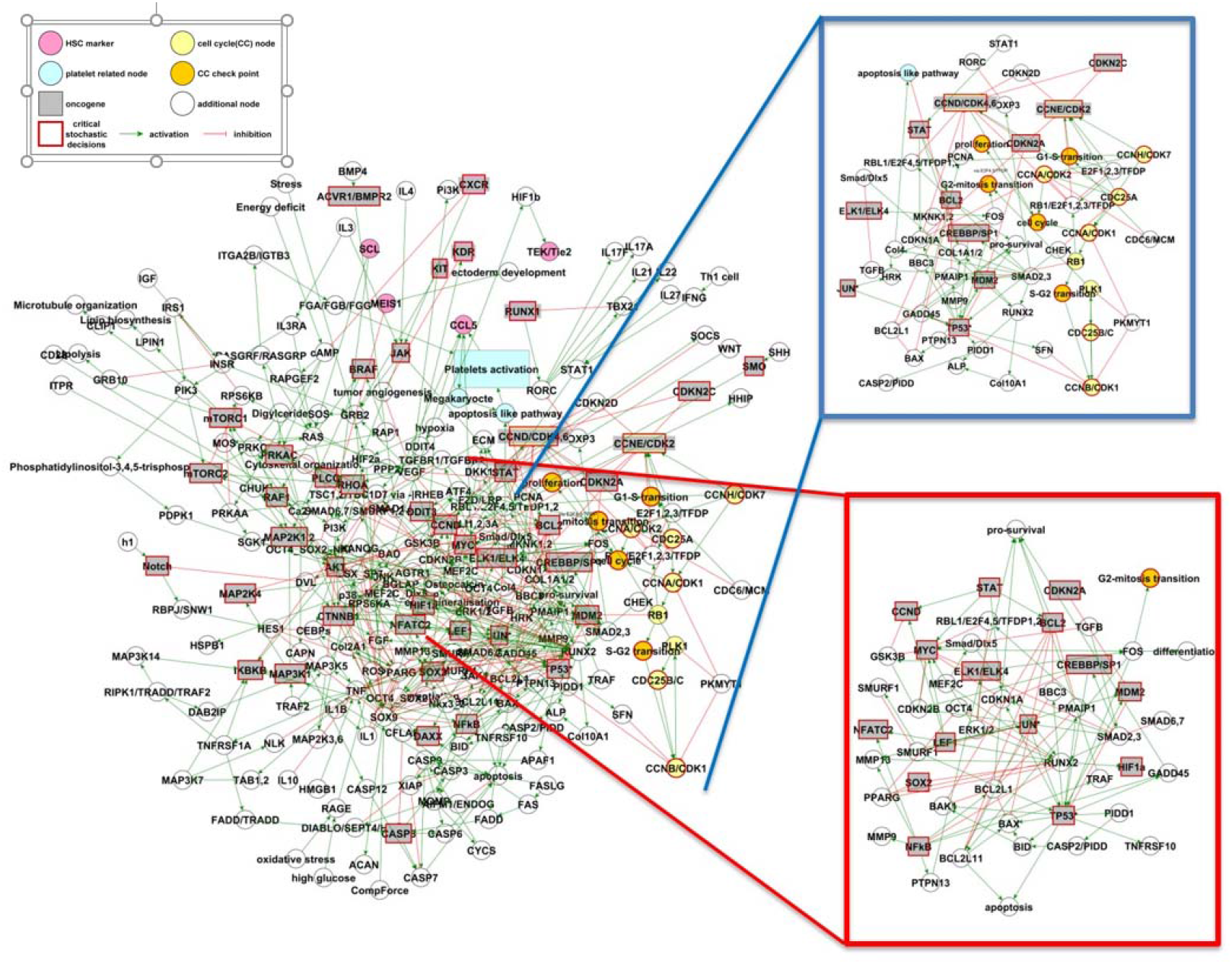
Full Hematopoietic Stem Cell (HSC) network. The network shown contains 242 nodes and 519 interactions. The inset in blue shows the cell cycle (top; Fig. 1b of the paper) and in red the cancer cycle (bottom; Fig. 1a of the paper). Parameters are the same as in Fig. 1a and b of the paper. Table S4 lists all proteins involved in the hematopoetic stem cells (HSC) network and related literature.

### 3. Full platelet network

**Figure S2:**
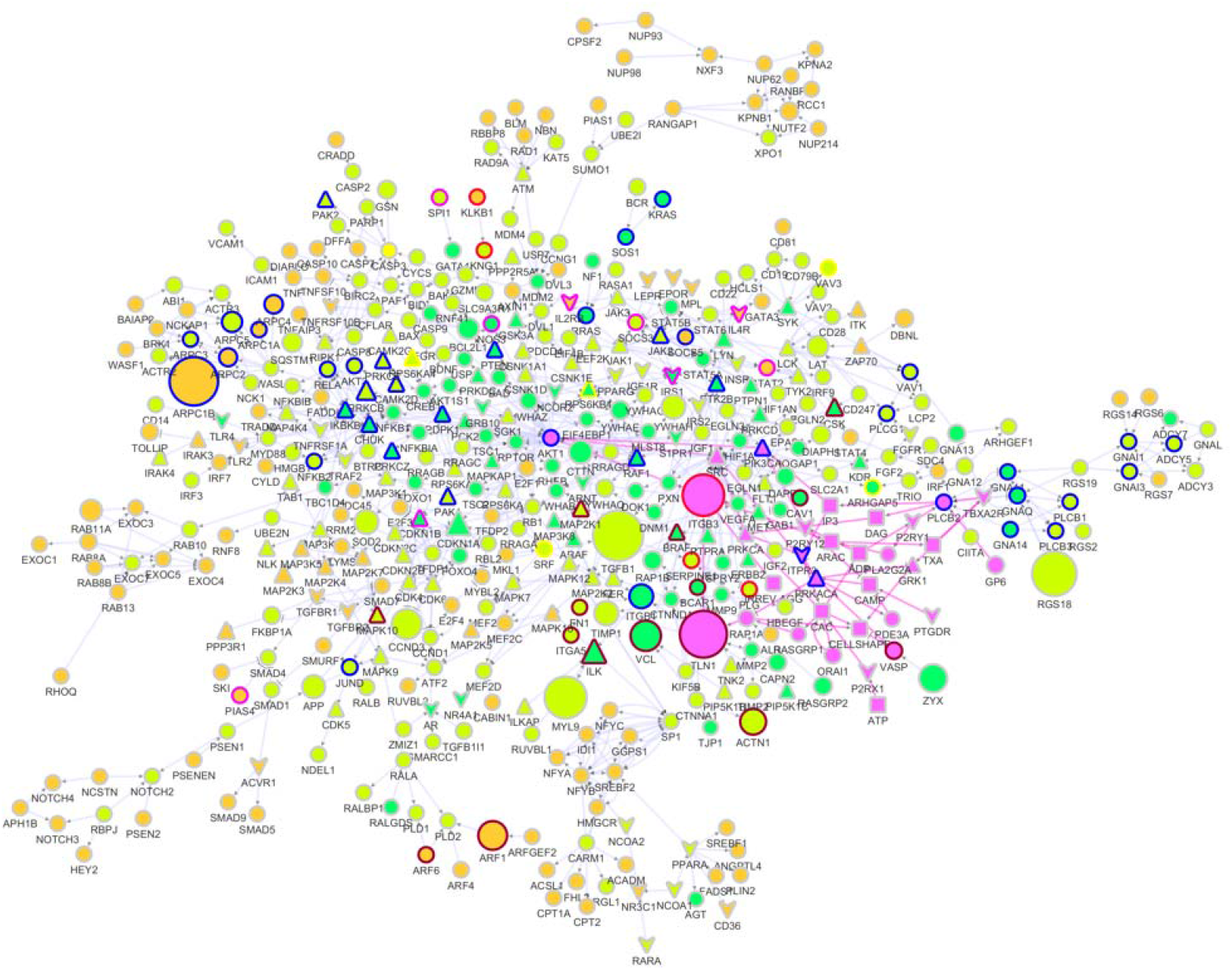
Full platelet decision network. This is shown up to neighbor degree 3 of the central cascade for activation and inhibition (455 nodes and 821 directed interactions). The full network contains as subnetwork the network shown in Figure 2 of the paper. Descriptions are alike, Fill color of nodes indicate the central cascade (purple), first neighbor (green), second neighbor (light green) and third neighbor (earth or orange). Calculations referring the centralities and stable states of the regulation of the central cascade where done using this network. Table S5 lists the platelet interactions of the central cascade (according to [5])

#### Unsure decision nodes

**Figure S3:**
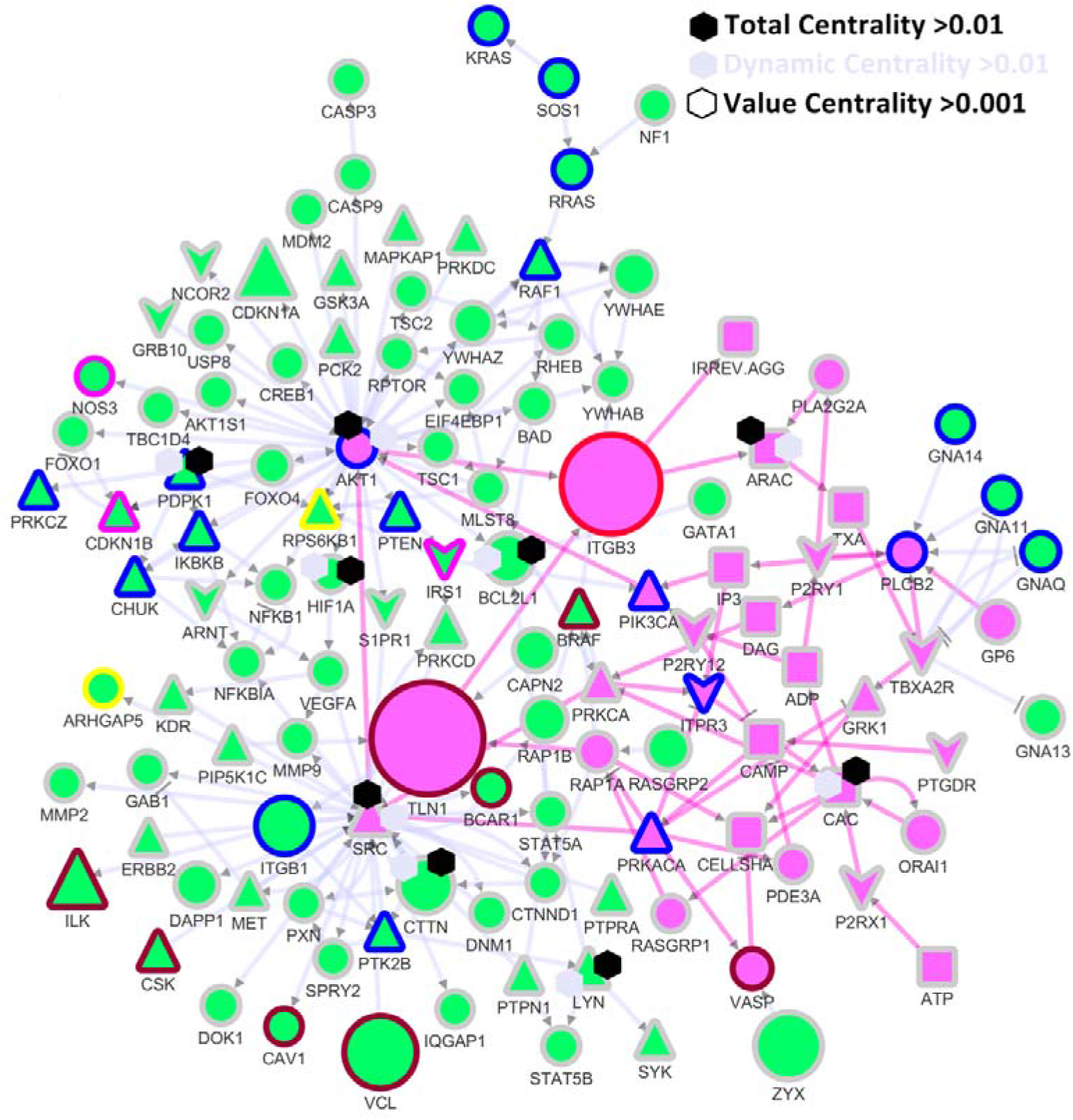
Central nodes in the platelet network. Nodes with high dynamic centrality are known to be key players in regulation, such as SRC, AKT1, cytosolic calcium (CAC) and ARAC. Compared are direct (VC), dynamic (DC), or combined network control (TC). The top nodes of Fig. 2b in any of these categories are labeled by hexagons. The purple fill indicates the central regulatory cascade. The green fill of the nodes represents the first neighbors. Further proteins: kinases, phosphatases (triangles), receptors (arrow head), macroscopic effects, non-protein coding molecules (rectangle) and protein-coding molecules (circle) are underlined. Pathways are indicated in border colors (red: blood coagulation; blue: inflammation mediated by chemokine and cytokine signaling; dark red: integrin signaling pathway; pink: interferon-gamma signaling, interleukin signaling and plasminogen activating cascade; yellow: PDGF signaling pathway). The node size represents expression in absolute values of RPKM.

### 4. Data on Cell division cycle

#### Decision processes not reachable for strict formal systems

Non-decidable statements or conflicting statements may arise from self-referring statements (for Gödel self-referring statements and “arithmoquination”, [6], see Box I in the paper). These are cellular processes states affecting and thus referring to the whole cell, e.g. cell cycle start is promoted according to dephosphorylation of cyclin B-bound cdc 2 for entry into mitosis. However, an inhibitory pathway (e.g. via wee1 catalysing the inhibitory tyrosine phosphorylation of CDC2/cyclin B kinase) may be activated at the same time: Internally then opposing information is not only represented but in the system opposing or non-decidable concatenations (for the internal rules how to represent information; Box G) from such statements can occur. However, as besides the internal representation each statement is also part of a molecular activity, answers happen nevertheless (in general stochastically) and can be e.g. apoptosis, recombination, proliferation etc according to the best survival chances in a potentially unlimited environment.

#### Cell division cycle decisions are complex

The number of all contained Kolmogorov processes K is compactly described by a program C (K) with length l. These are all processes to be represented by a shorter program providing the same output. Next we collect all non-compressible phenomena from the type of O(N) numbers (Chaitin complexity[7]). In particular, process averages, stopping probabilities etc. are non-compressible complex. Moreover, there are more complex interacting non-compressible phenomena such as self-organising processes and life (O(network) numbers e.g. species, DNAs, etc.) based on decision process networks:

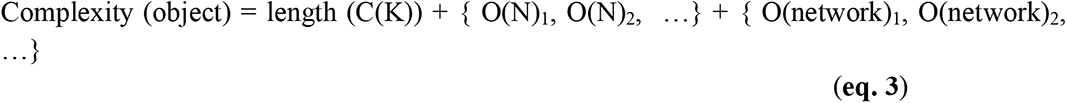

In total, we have a rather complex decision complex as the resulting program is neither linear nor compressible. This is typical for decision processes and illustrated now in concrete detail looking at the cell cycle:

(a) Cell Cycle Reactions

**Fig. S4.**
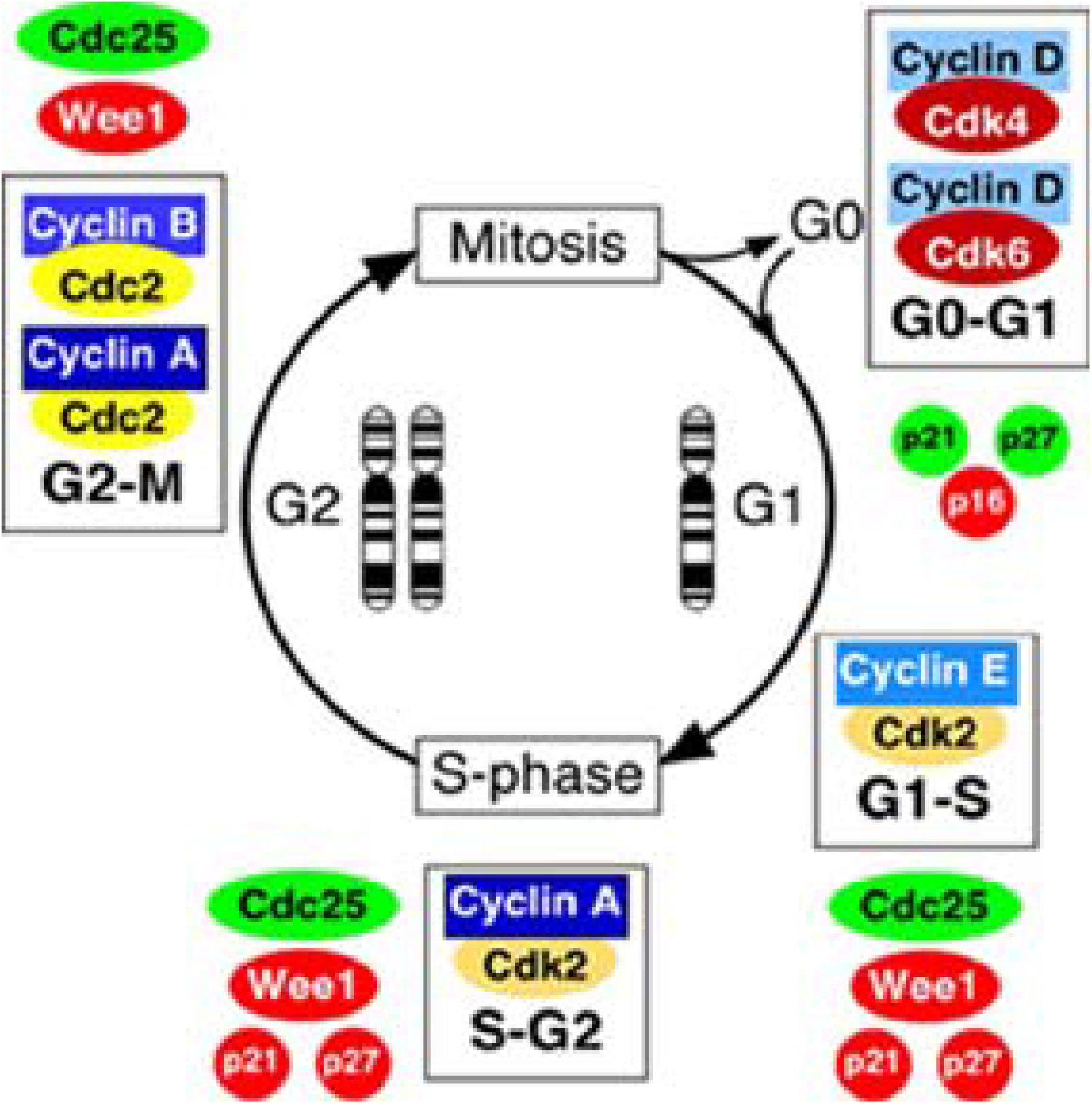
Cell Cycle Reactions. The cell cycle ensures duplication of the DNA and Chromosomes (central image). Shown further are critical phosphorylation and dephosphorylation steps, in particular of Wee1 protein (red ellipse) at position 21 and 27 (red cycle) and the interaction with the different cyclin complexes (grey square including cell cycle proteins cdk4 and 6, top right, shown again as red ellipses) and how this relates to the different cell cycle transitions (bottom line in each square). We next list in detail the specific reactions for a typical eukaryotic cell according to the reactome map.

**(b) Cell Cycle steps (see also Reactome map): Sub-map G to S transition**

~~~
c Cyclin E associated events during G1/S transition
  c Formation of Cyclin E:Cdk2 complexes
   m Formation of Cyclin E1:Cdk2 complexes
   m Formation of Cyclin E2:Cdk2 complexes
  c Translocation of Cyclin E:Cdk2 complex to the nucleus
   m Translocation of cyclin E1:Cdk2 complexes to the nucleus
   m Translocation of cyclin E2:Cdk2 complexes to the nucleus
  c Inactivation of Cyclin E:Cdk2 complexes by p27/p21
   m Inactivation of Cyclin E1:Cdk2 complex by p27/p21
   m Inactivation of Cyclin E2:Cdk2 complex by p27/p21
  c SCF(Skp2)-mediated degradation of p27/p21
  c Cyclin E/A:Cdk2-mediated phosphorylation of p27/p21
   m Cyclin A:Cdk2 mediated phosphorylation of p27/p21
   m Cyclin E:Cdk2-mediated phosphorylation of p27/p21
  c Association of Cks1 with SCF(Skp2) complex
  c Binding of phospho-p27/p21:Cdk2:Cyclin E/A to the SCF(Skp2):Cks1 complex
  c Ubiquitination of phospho-p27/p21
  c Degradation of ubiquitinated p27/p21 by the 26S proteasome
  c Phosphorylation of Cyclin E:Cdk2 complexes
   m Phosphorylation of Cyclin E1:Cdk2 complexes by Wee1
   m Phosphorylation of Cyclin E2:Cdk2 complexes by Wee1
  c Dephosphorylation of Cyclin E:Cdk2 complexes by Cdc25A
   m Dephosphorylation of Cyclin E1:Cdk2 complexes by Cdc25A
   m Dephosphorylation of Cyclin E2:Cdk2 complexes by Cdc25A
  c CAK-mediated phosphorylation of Cyclin E:Cdk2
  c Phosphorylation of proteins involved in G1/S transition by active CyclinE:Cdk2 complexes
  c Cyclin E:Cdk2-mediated phosphorylation of Rb
 c Ubiquitin-Dependent Degradation of Cyclin E
  c Ubiquitin-Dependent Degradation of Cyclin E1
  c Ubiquitin-Dependent Degradation of Cyclin E2
 c G1/S-Specific Transcription…
~~~

#### Complexity measures examined for cell cycle decision processes

The cell cycle (**Fig. S4**) can then be examined in detail regarding the complexity levels of the involved decision processes. By this, different levels of complexity involved become clear Using the software cytoscape, the different complexity measures are exemplified for Reactome maps ([8]; http://www.genomeknowledge.org:8000/about.html) of the cell cycle.

**(a)** Shown is the central biological circuit for the G1 to S transition (in colour different involved proteins). **(b)** The simplified Reactome map for the cell cycle examined here contains 356 molecules in the simplified cell cycle model of the Reactome database (map React_152.2). The biochemical key events of decision events are given as a marked list.

**Kolmogorov compression** endeavours to represent the behaviour of this system by a shorter programme giving the desired output (e.g. replicate / do not replicate; thus shortest conceivable but non-predictive trivial model contains one bit), trivial compressions are all representations of the cell cycle using fewer nodes (though the output characteristics would not be identical). Instead the number of genes involved in the cell cycle involves hundreds of genes ([9]; the Kolmogorov complexity in this more accurate model is already in the order of tens of thousands of bits).

Moreover, we show that *no short programme* can represent the resulting complexity for the cellular *transition processes* involved, e.g. the transition from G phase to S phase (this is more complex then Kolmogorov, the Chaitin complexity, see [7] participating are 127 molecules, if the network is set up according to the genomeknowledge database. A key interaction is between cyclin E and cdk 2. The programme describing this level of complexity is at least as long as the information content of the protein sequences and the regulatory sequences of the genes involved (at least in the order of hundreds to thousands of bits). The non-compressibility can also be shown genetically: All information carrying parts of the system do in fact change the behaviour in case of mutation.

Including interactions renders the complexity even higher (O(network)). This is in principle unlimitedly more complex if all details of interactions are considered including environmental interactions and exact behaviour in time for the cell division programme under all conditions. For example, data on cell division kinase 1 phosphorylation targets involve over 500 proteins in yeast[10] some of them participate in feedback regulation.

Interestingly, simulating such cellular conditions (e.g. using Cytoscape, see **Fig. S4** or Jimena, **Fig. S1-S3**) referring to the whole cell and involving large genetic circuits shows that opposing signals can often lead “to blocks” of the system (e.g. not only regarding cell cycle but also decisions such as apoptosis vs proliferation). However, as besides the internal information there is also the actual enzymatic activity encoded from the circuit, a clear behaviour arises in most cases in a finite time with a clear preference for one state (sometimes including a “blocked” state but then also with biological meaning and selection advantage, e.g. G_0_ phase in differentiated cells). Note furthermore that the system behaviour on the phenotypic side (so beyond the internal representation capabilities of the system) is selected by the environment (itself part of a potential infinite context) for the benefit and best survival capabilities of the organism. Thus “crossing the Gödel limit” often happens in living beings in complex decisions affecting the survival of the whole cell or higher organism and regarding the decision between different survival strategies.

Examples include apoptosis versus proliferation, recombination or not as well as proliferation or differentiation. The identification of the best survival strategy or cell cycle replication (Figure 3) strategy at a given environmental condition is in fact a non-decidable Gödel problem for a formal system considering the complexity and stochasticity of the transitions and interactions involved and the potential unlimited environment. The calculation of mutation effects in regulatory proteins for fitness and survival, e.g. mutations in cdk2 is similarly challenging. There are further levels of complexity added in multi-cellular and neural organisms. Detailed calculation and application examples include e.g. the decision of RIP protein between apoptosis or replication where details of the interaction surfaces and the sequence of protein encounters determine the resulting decision [11], or the triggering of a differentiation signal versus malignant proliferation where concentrations and ratios of two different isoforms of Raf, B-Raf and C-Raf, determine cellular fate. Stochastic processes in the cell result from this. In general they describe the complex cellular adaptive behaviour better than non-stochastic models (e.g for the important cell cycle transition from G1 to S, Reactome map <u>REACT_1783.2</u>). The same reasoning applies to complex decisions by higher organisms as well as regarding machine-implemented complex decision processes (including nano-machines).

These arguments explain why fundamental cellular processes such as differentiation, cancer, apoptosis, aging and inflammation are highly complex (O(network) processes – more complex then computer programs, see above) and inherently undetermined (e.g. cancer, aging). Hence only modelling as stochastic process is accurate and detailed modelling is only possible by iterative refinement (including experimental data and data-driven parameter fitting and modelling). Own examples include platelet activation and inhibition [12], cardiac hypertrophy [13], bacterial growth including metabolism [14], plant networks and their regulation [15], as well as animal regulatory networks [16]. Oncogenes such as c-Myc exemplify this (combining stochastic events leading to cancer mutations with general transcriptional enhancer as well as specific DNA binding and protein complex formation) as well as aging as a highly complex cellular process (combining again different stochastic aging processes and determining cellular programs combine here).

### 5. Further analysis of cellular decision processes

#### 5.1. Comparing decision processes

Pinpointing biological heuristics, identifying involved decision nodes and processes in platelet and hematopoetic stem cells elucidates inherent risks for cancer and cardiovascular diseases and gives suggestions for improved search strategies dealing with NP problems.

Specifically, we show in molecular detail how Gödel decision dilemmas arise and also how they are avoided in biological systems by stochastic resolution of the decision conflict. We show that these arise in the cell cycle and in accordance with this there should be a linear increase in the risk of errors (cancer) with the number of cell divisions as described recently[3, 4]. We pinpoint this to self-referring statements and extend this to oncogenes. Large-scale networks and simulations (from 100 decision nodes to up to 5500 nodes) on platelets as well as their stem cells megakaryocytes show here for the first time different types of decision nodes (stable, unstable, undecided; direct, dynamic or total network control) and how in the platelet the central activating cascade, inhibition and cytoskeleton is involved. Each node and all networks simulated were carefully validated (see supplement) and we see that in addition to the cancer risk by cell division specific oncogenes participate in the hematopoetic stem cell (Fig.2; name oncogenes) for this type of error-prone system instability instead of healthy and controlled cell division. Regarding thrombus formation and the two major killers stroke and heart attack we accumulate here evidence how and in which proteins the delicate balance between blood flow and blood stasis can be easily tripped by unstable decision nodes and platelet crosstalk including new and additional points for pharmacological intervention influencing the involved proteins.

We show thus that in biology heuristics prevents too long decision processes and this can be applied to solve NP problems such as protein folding. A maxcut solution strategy achieves close to optimal solutions [17]. We used such a strategy as new protein fold prediction strategy. This was tested by us in 2D systems but something similar happens in real protein folding for the nascent chain from the ribosome [18]. In the latest CASP competitions a key player in the accuracy improvement, in the free model scheme of protein folding, is the contact prediction [19]. We explore the possibility of using the max-cut algorithm to make the contact predictions. We also show that protein folding is starting conformation dependent, a bad starting conformation leads to aberrant protein structures and plaques formation.

Similar concepts have been proposed as folding funnels and to explain neurodegenerative disease, but here we show how all this derives directly from our comparative study of decision processes in biology. For cellular networks, NP problems involve again decision processes, and our platelet and HSC simulations are only possible applying heuristics.

Finally, we think that further complexity comes into NP problems by ambiguous possibilities for solution representation and illustrate this by an example (Fig. 4b). The “given sum problem” is a related problem with proven NP properties [20]. Computers definitely may take much longer than expected to compare solutions among multiple representations for transcendent numbers (no algebraic polynom solutions; e.g. Lindemann-Weierstrass theorem; [21]) whereas human beings have in everyday life to come up with solutions in finite time and do not encounter a halting problem at fundamental decisions such as Gödel incompleteness.

Another way to phrase the capabilities of living observers is that they have a concept of meaning and can hence organize decision processes efficiently for survival though the environment is complex and potentially infinite. In this sense “we have an understanding of concept of infinity” [22] which a machine does not share and hence cannot cope with. Mathematics clearly has a notion of infinite. In our everyday world, smallest components and a limited observation horizon lead to a finite environment. however, every environment is part of a potentially infinite complex world including sudden mutation by radiation from distant as well as the infinite possibilities for quantum states, virtual particles, and Feynman path integrals.

#### 5.2. SAT equivalence and implications

Cell signaling can be stated as a SAT problem. We show that it is even equivalent to a SAT problem (see p. 26 in this supplement). It is hence **NP** complete, too and this is also true for metabolism and metabolic fluxes (here the “combinatorial explosion” for calculating all possible modes, so being confronted with an NP problem, is well known). Again, cell signaling is not perfectly accurately executed, but rather a heuristic solution guarantees that on average the reaction given ensures maximum survival probability (otherwise it would not have been selected that way). In metabolism there are heuristic solutions (in particular metabolic channeling, compartments, major pathways and their regulation) probed quite often. However, the complete metabolism has an inbuilt parallelism of all metabolites being operated in the cell in parallel, so for the full and detailed metabolic equilibrium it can again be argued that here the NP problem is solved in almost full detail exploiting clever heuristics (in particular regarding metabolic regulation, pace maker enzymes and allosteric regulation) but also parallel calculations, by all metabolites as involved molecule numbers are high enough and affinities and enzymes operate sufficiently fast and direct. Regarding cellular networks, fast system decisions rely on an optimized small network of key nodes including modulatory cross-talk. Semi-quantitative Boolean modeling such as cell differentiation of heart insufficiency or platelet activation is surprisingly powerful to model cellular decision networks. Such predictions agree well with experiments after several iterative cycles of theory, refined prediction and refined experiment. Evolution tackles NP problems such as optimal protein structures or phenotypes for survival in general really head on: it takes non-deterministic polynomial time to identify solutions in numerous parallel trials in many genomes over long time.

However, regarding physics, limitations of biology sometimes may play no role, for instance in quantum physics, hence here more demanding computations are possible. We are aware that even a quantum computer may not allow to solve all NP problems: it efficiently solves factoring and discrete logarithm calculations, however, graph isomorphisms and NP complete problems may still pose a challenge ([23]; illustrated in Fig. 3 of the paper). However, and in contrast, natural physical processes really emulate all combinatorial events and paths in parallel, for instance the Feynman path integral traces all energy levels and trajectories. Only by this massive inbuilt quantum parallelism we are able to find the observed trajectory and so here the complete problem space is covered for quantum processes.

#### 5.3. NP problems are tough to simplify and heuristically yet error prone solved in cells

Regarding mathematics, originally we had in mind to proof that we can rule out some NP problems as ill-defined, and, using this knowledge, we solve the remaining ones in polynomial time. However, something more complex seems to be the case: There is higher complexity, sometimes non-obvious in NP problems preventing their simplification or solution by a well-defined solution strategy with polynomial calculation time:

All NP problems including the Boolean satisfiability problem (SAT) are equivalent (Cook’s theorem; [1, 2]). If you could solve a SAT problem in **P** you would also be able to solve all **NP** problems in **P** applying the theorem of Cook.

Where is now the opacity that prevents simplification of search strategies in NP problems? To pinpoint this, we start with a toy example (Fig. 3 in results): Consider a program which determines if two numbers are equal or not (given as decimal digits). If you take rational finite numbers, you check digit for digit whether each is equal and once you reached the last digit you are done. The calculation time grows linear with the number of digits to check (P problem). If you consider instead real numbers there are a lot more numbers (cardinality is no longer countable infinite and hence much, much more). Now you can easily verify that two numbers are different if shown at least one example where the digits are different. However, now there is a combinatorial aspect in the comparison problem included. To find the shortest description for real numbers is, in our opinion, equivalent to the subset sum problem. Regarding mathematical implications we think this corresponds to constructivism (rejection of actual infinity). We suggest that for modelling the many fast heuristics and decision processes in biology, it is better to choose a model where the continuum hypothesis is not used and there are no intermediate cardinalities of infinity. In living cells, a heuristic and fast cellular decision process stops deterministic in polynomial time with the risk of occasional errors regarding the beneficial effects of the decision. Such a deterministic process corresponds to a clear description of a number in our toy example. For the description of such clear deterministic processes, a subset of **R** (real numbers) is sufficient. It has the same cardinality as **Q** (rational numbers) or **N** (natural numbers) and contains in addition only all irrational numbers that have a clear description. However, everything more complex are problems producing undetermined halting behavior and there is then also no mapping of these higher cardinality sets of numbers to **Q** or **N**. We suggest this looking at a toy problem, comparing two numbers. However, we think that looking for a shortest **and unique description** of a number is equivalent to finding the shortest program for its correct calculation and that one can show that this problem allowing all numbers including sets of cardinality **R** is SAT equivalent and hence, that really any problem that has in this sense no longer a simple **or unique description** and is in this sense clearly more complex is at the root of the challenge of NP problems.

This is suggested from biology and definitely valid for biological systems. However, what this exactly means in mathematical terms has still to be found out.

In conclusion, pinpointing biological heuristics, identifying involved decision nodes and processes in platelet and hematopoietic stem cells elucidates inherent risks for cancer and cardiovascular diseases and gives suggestions for improved search strategies including suggestions for NP problems.

#### 5.4. Medical and evolutionary implications

Interestingly, diagnosis of heart attack is exactly for this reason relying on diagnosing an unstable thrombus. However, this instability cannot be recognized by nuclear magnetic resonance (NMR), ultrasound, or other means. Hence, we can at least pinpoint here the roots of this medically highly important problem (Fig. 2 in results). **Table S1** indicates experimental data (gene and protein expression, gene knockout of GPVIb against wild type) and network analysis of these validation data. Any new decision points revealed can now easily be targeted by drugs, our resource, the PlateletWeb (http://plateletweb.bioapps.biozentrum.uni-wuerzburg.de/plateletweb.php), gives many suggestions including already clinical approved drugs.

Similarly, for a model of the hematopoietic stem cell differentiation (Fig. 1) we show that also fundamental cell cycle decisions are stochastically resolved, lead to megakaryocyte differentiation and platelet formation. A linear increased risk of decision errors implies a stochastic activation of oncogenes for cancers of lymphoma and leukemia type.

Biology is not perfect as population size is never infinite and rather limited for many members of the megafauna. Evolutionary paths are limited by fitness gain and neutral paths theory. Only typical large bacterial populations solve truly optimization problems in parallel and with high success including genetic adaptation to NP problems such as optimal resistant protein structures against new antibiotics.

### 6. Protein folding and cellular signaling as decision problems

Biological Heuristics simplify NP problems to be solvable in P deriving pareto-optimal solutions leading to a new protein folding heuristics by constraint based optimization and proof that cellular decisions are equivalent to SAT problems.

#### 6.1. On Nature solving NP problems

We believe nature does not solve **NP** problems in biology. This statement cannot be demonstrated rigorously but we present several different cases occurring in nature where it can be seen that nature solved a relaxed version of the respective **NP** problem, taking always a pragmatic approach. We study two main cases, protein folding and cell signaling networks. We also present R**N**A folding and cell metabolism as analogous cases to each of the principal cases.

Regarding protein folding, evolution has almost “infinite” time, billions of individuals and positive epistasis with clear paths. Furthermore, at least physics “knows” always the correct solution according to the law of energy, so does this “solve” an **NP** complete problem?

Or a bit more general, looking at bioinformatics, you anyway reduce your **NP** problem to a **P** problem. Thus RNA folding can be simplified for just two dimension or secondary structure predictions in a homology model ignoring loop regions: All are **P** (quadratic, cubic and so on). 3D folding sometimes is also reduced to **P** problems using Boolean decisions, system state enumeration etc. However, all these natural occurring problems with combinatorial flavor are from the accurate mathematics **NP** problems.

#### 6.2. Protein folding (see Fig. 3a)

The protein folding problem arises from the accepted believe that the tertiary structure or native conformation of the protein is encoded in its primary structure or amino acid chain, at least for small globular proteins this is fully proven to be true. The protein folding problem consists in predicting the native conformation based on the amino sequence. Several models have been proposed for the problem and most of them have a clear proof to be **NP**-complete, for example the hydrophobic model has a reduction to Bin Packing problem and Protein Threading was also proven to be **NP**-complete. These models are abstractions from the real problem as seen in nature so it can be argued that they do not really say anything about the complexity of the real problem. For this reason we present a theoretical model which more clearly resembles the real protein folding problem as seen in nature and show this way that the problem is **NP**-complete. Based on this model we propose that nature does not solve here the **NP**-complete problem but a relaxed version of it.

##### 6.2.1. Protein Folding as a Maxcut problem

On this model, the amino acid sequence is modeled as a graph, where each amino acid is a vertex and each weighted edge encodes the value of the free energy reduction during the folding due to the two amino acids linked by that edge.

**Figure S5:**
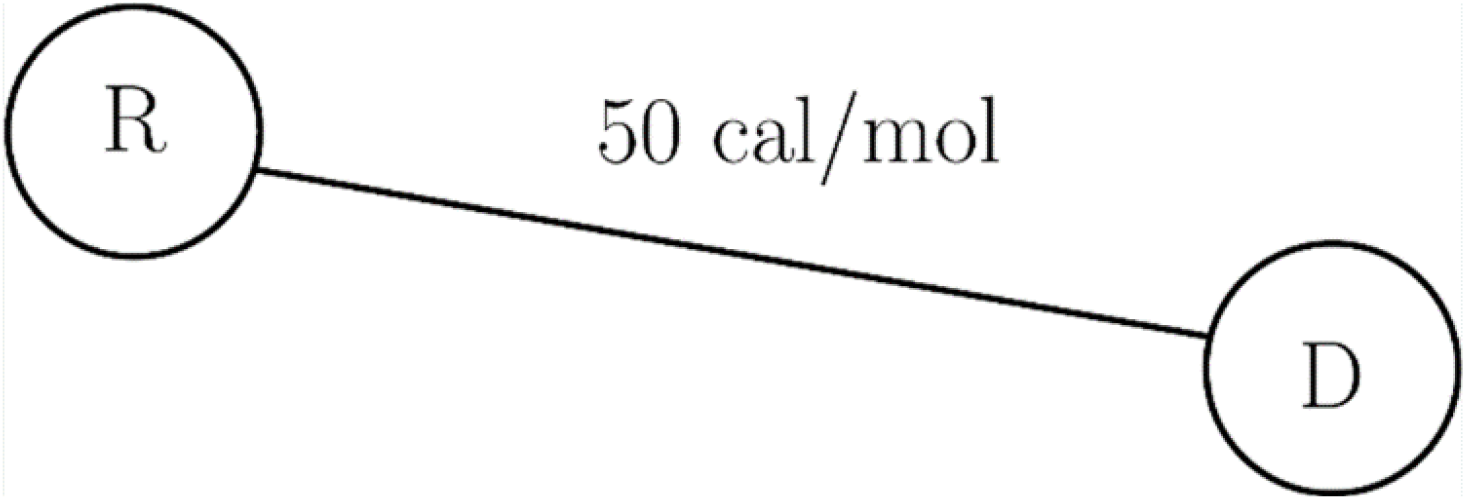
Proteinfolding represented as a MaxCut Problem. To look at protein folding form this angle, Amino acid representation is done as vertexes and the interaction represented by the vertex. R for Arginine, D for Aspartic acid and a hypothetical interaction of 50 cal/mol.

Based on this representation the primary structure of the protein can be presented as a full graph. This graph has possibly edges.

Given a graph where is the set of vertices and the set of edges, the Maxcut problem consists in splitting the graph in two sub-graphs, so each vertex belongs to one of the two sub-graphs, while assuring that the splitting passes through the edge combination with the highest weight.

**Figure S6:**
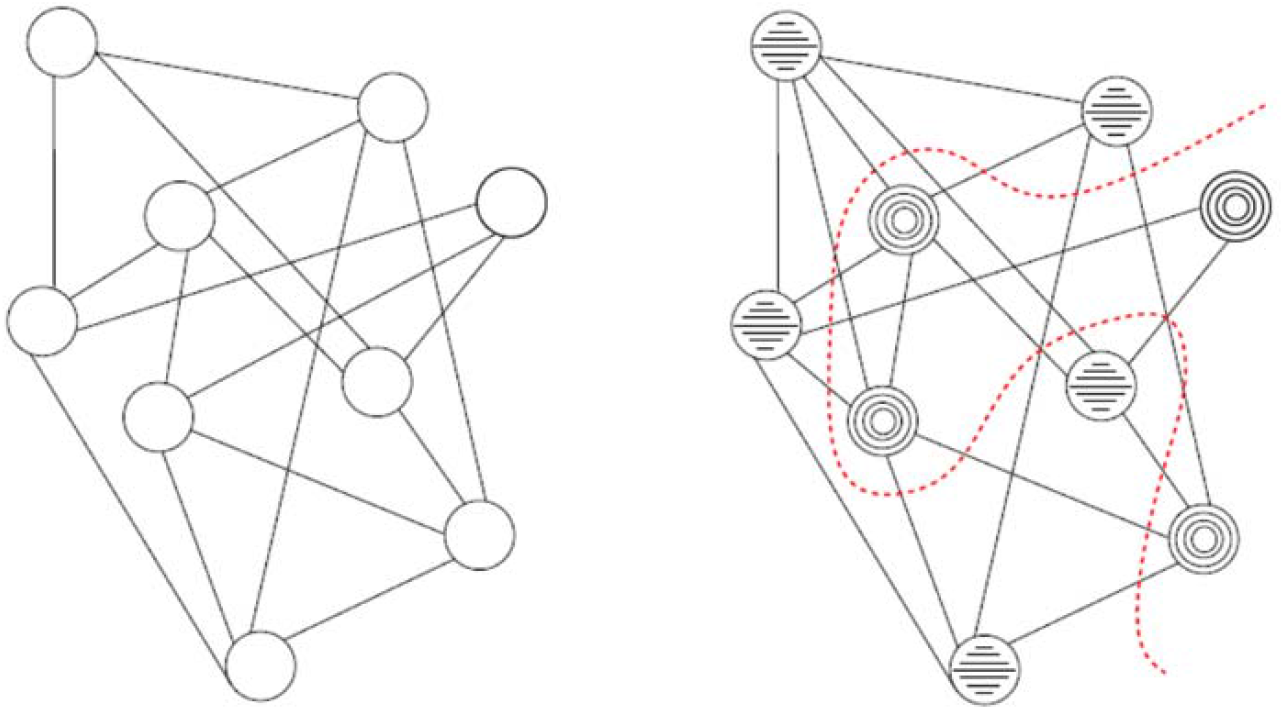
Example protein represented as a MaxCut Problem. An example graph represents now a small protein and its Maxcut solution is shown.

By stating the protein folding model like this we see that the Maxcut solution corresponds to find the set of edges that account for the bigger drop in the free energy. This model is a recursive continuous model, on each step the interactions are recalculated and the Maxcut algorithm states the most favorable route to follow. The model is theoretical for two reasons, the first, that calculating all the interactions between the amino acids is if not impossible at least at hard as the protein folding problem itself and the second reason is because the Maxcut problem is **NP**-complete so it cannot be efficiently solved.

The first obstacle seems unsolvable but the second one is not. Using the Goemans and Williamson (2004) algorithm[17] the Maxcut problem can be solved with at least a 0.878 approximation. We believe that nature actually boards this approach while folding proteins. This believe comes from the fact that the amino acid chain is not a rigid code for the tertiary structure but on the opposite it accepts a lot of changes while still folding to the same structure, this is a form of relaxation like the one observed in the semidefinite programming step[24] of the Goemans Williamson algorithm. Additionally, we see that the algorithm makes use of randomness and in nature as well has been seen that the proteins experiment a lot of random movements and vibrations that allow to search for the proper conformation. This same logic applies to the RNA folding problem including simplified versions such as RNA secondary structure.

The following code is an script to solve the 2D lattice problem of protein folding based on a simple convex optimization problem. Given the initial conditions protein folding is just a search for a local minimum. In this example a small globular protein is used, the protein is 46 aminoacids long. In order to use the code the users have to install R and the Gurobi optimization package.

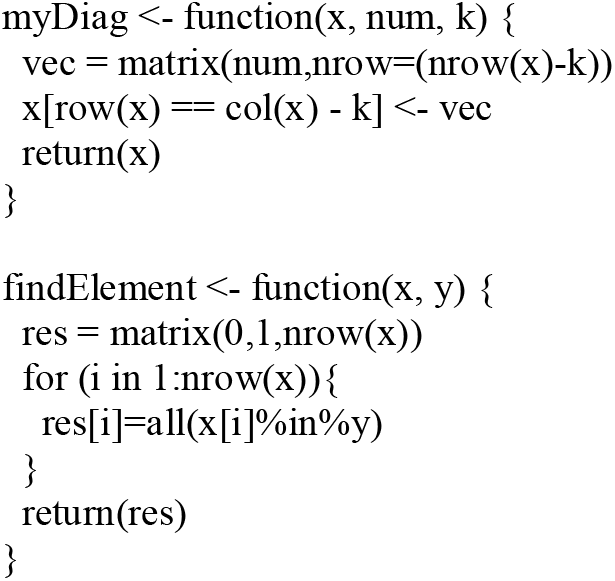
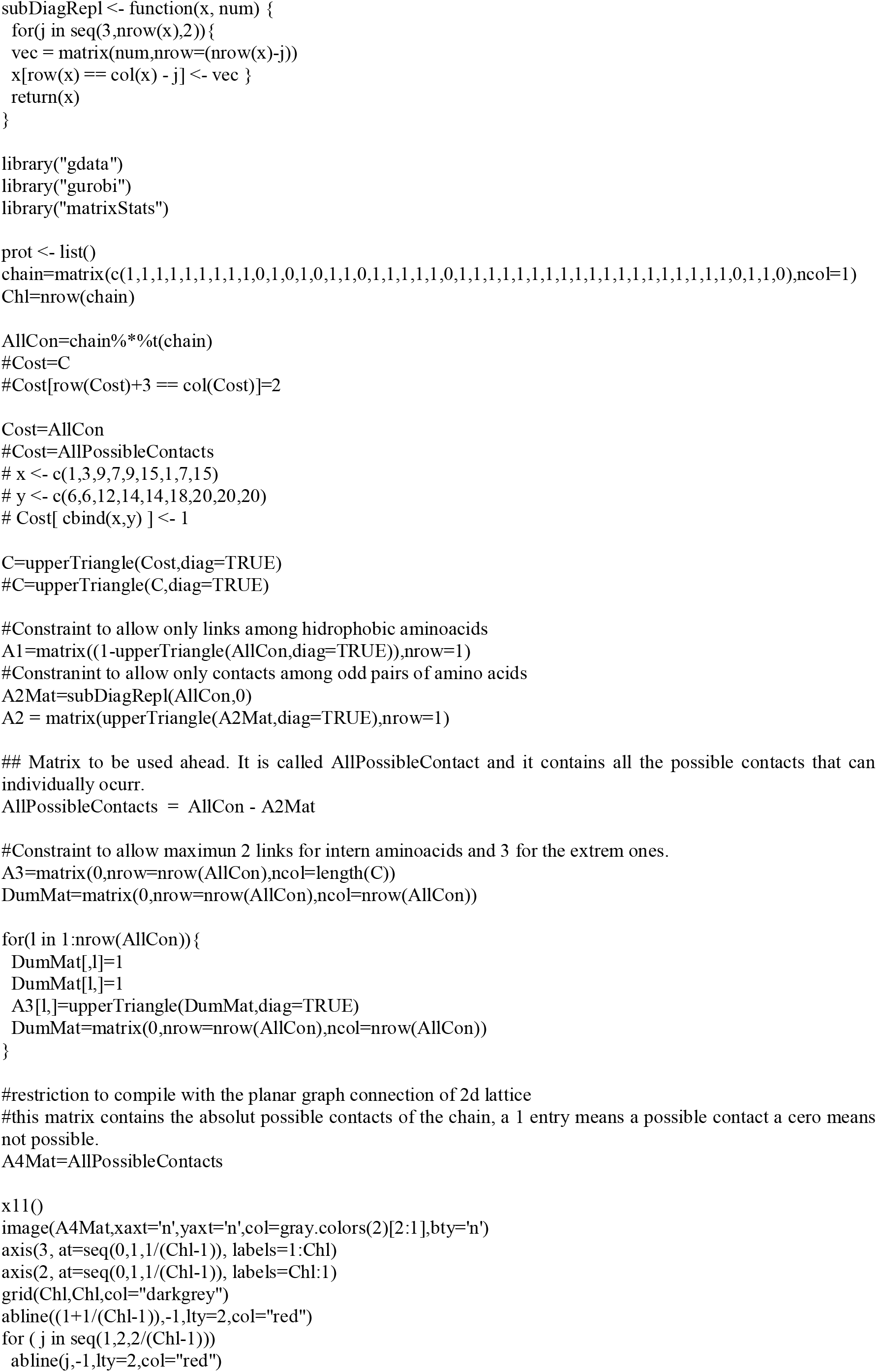
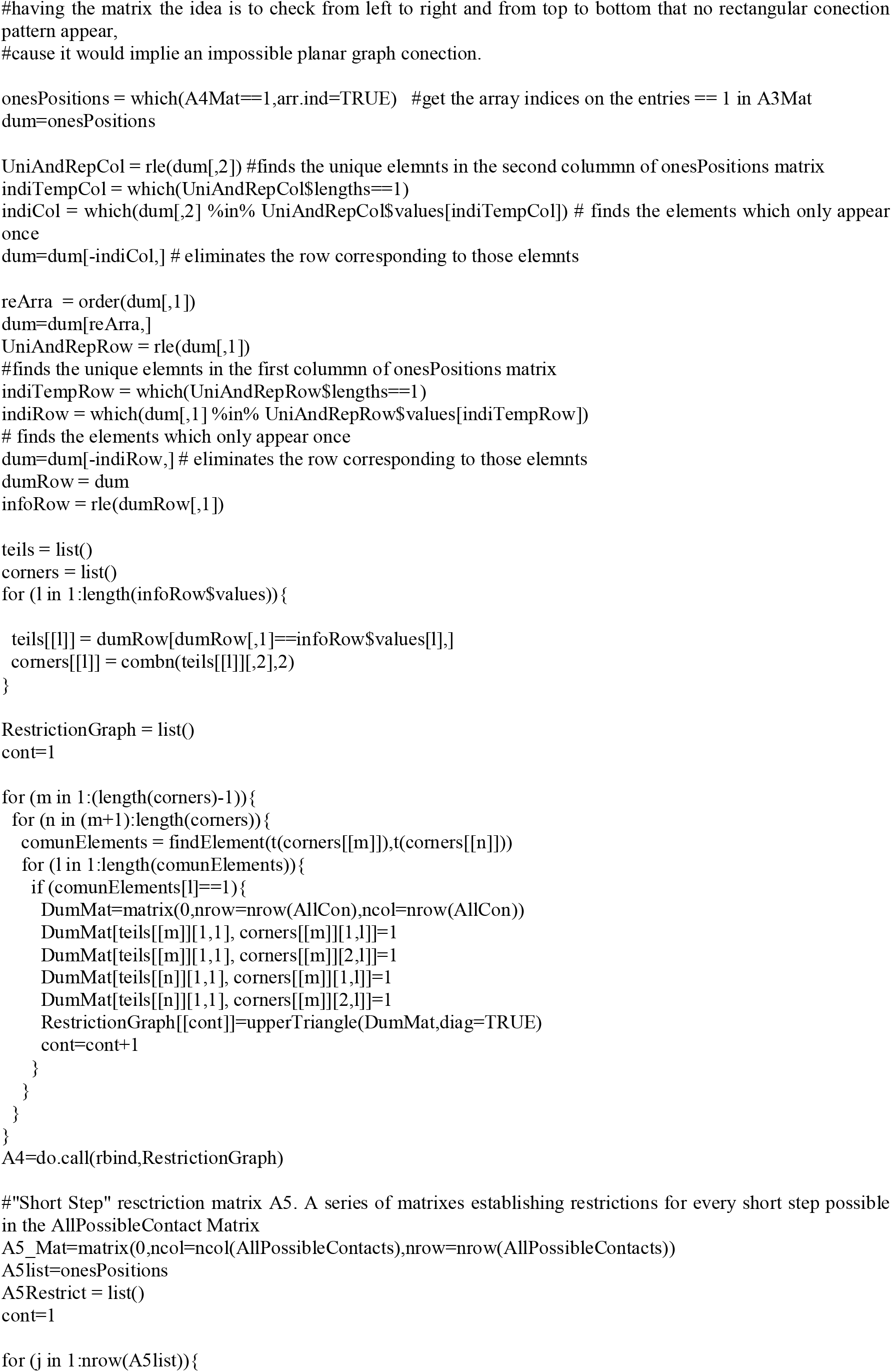
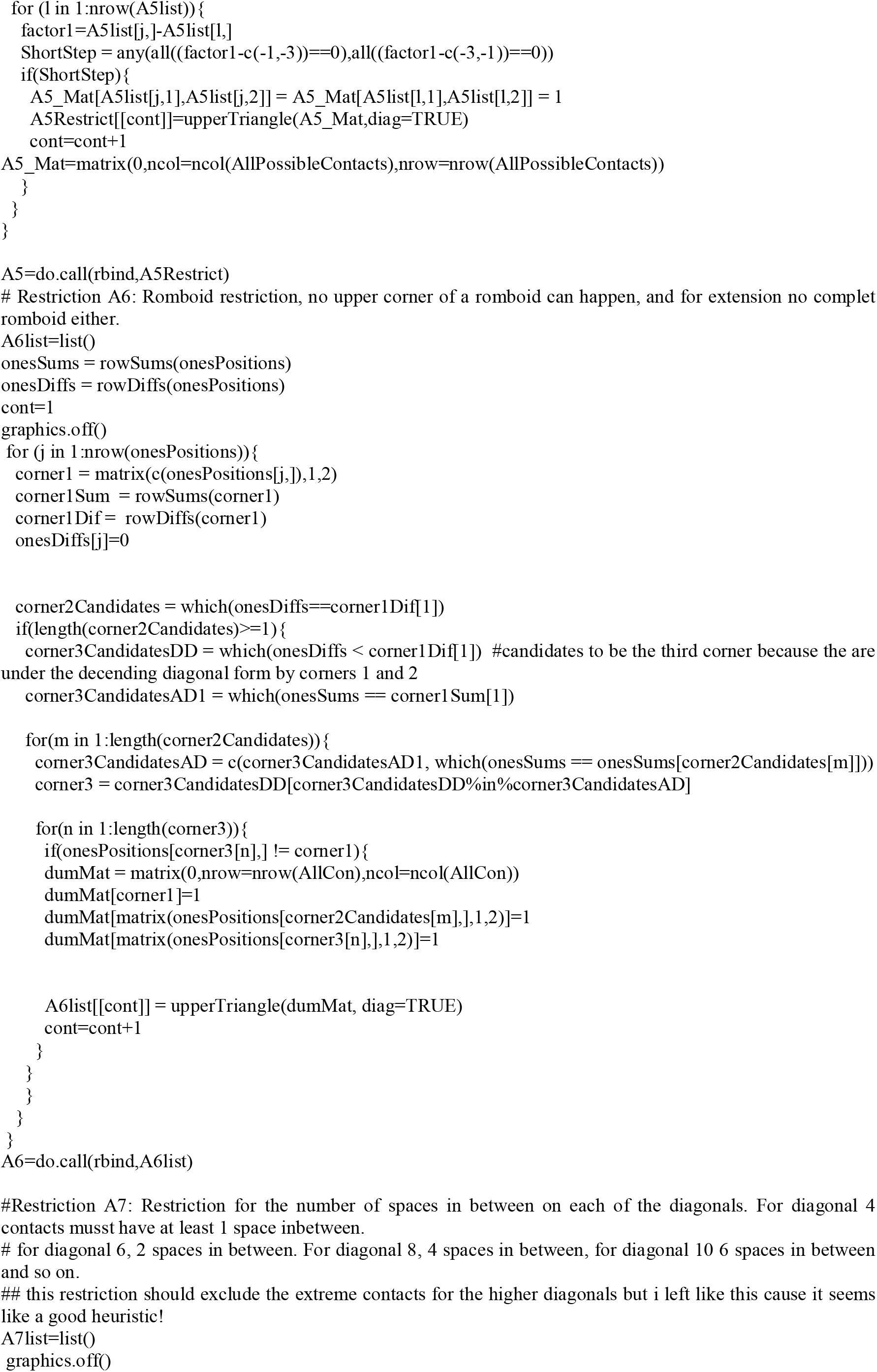
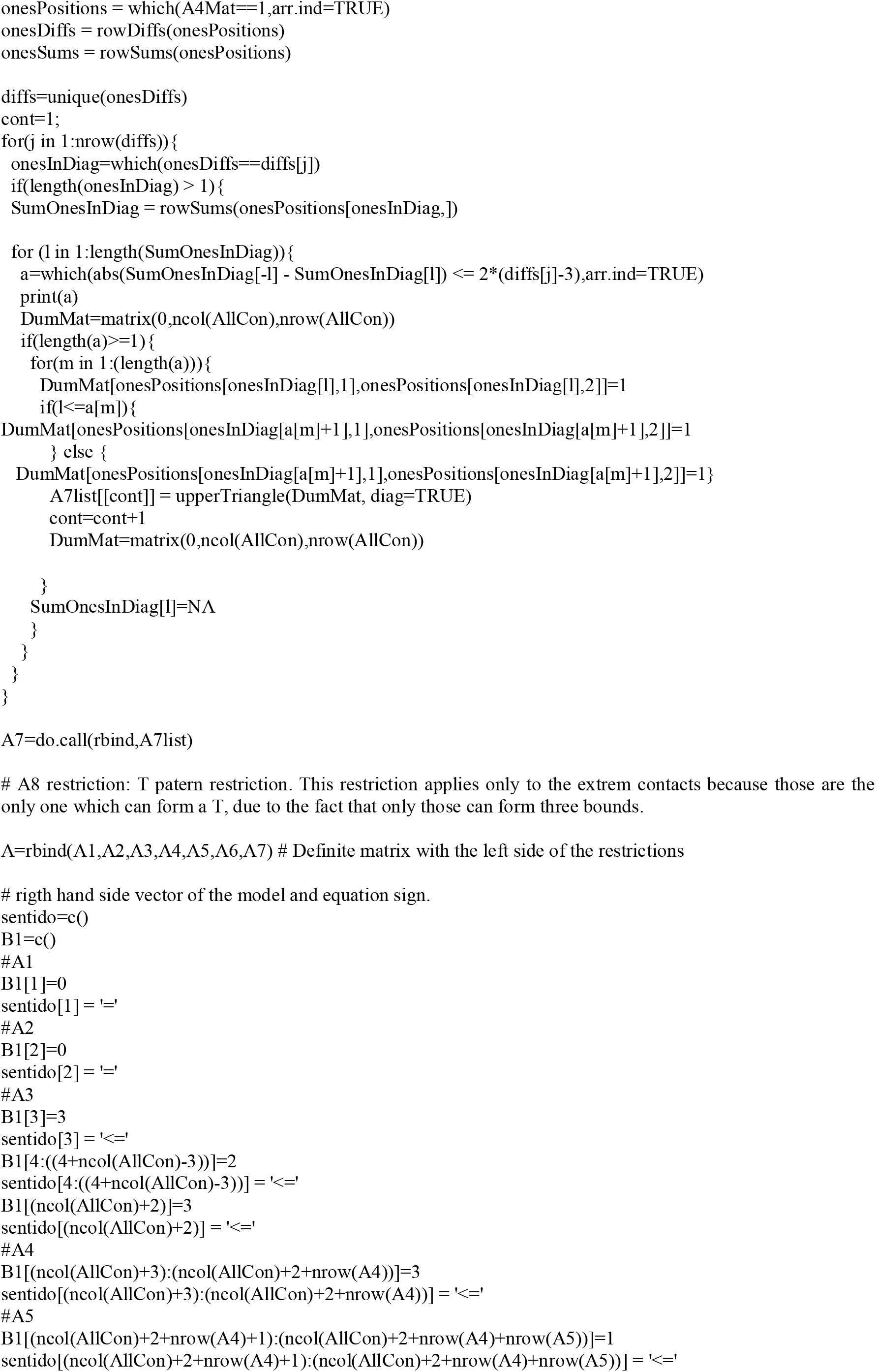
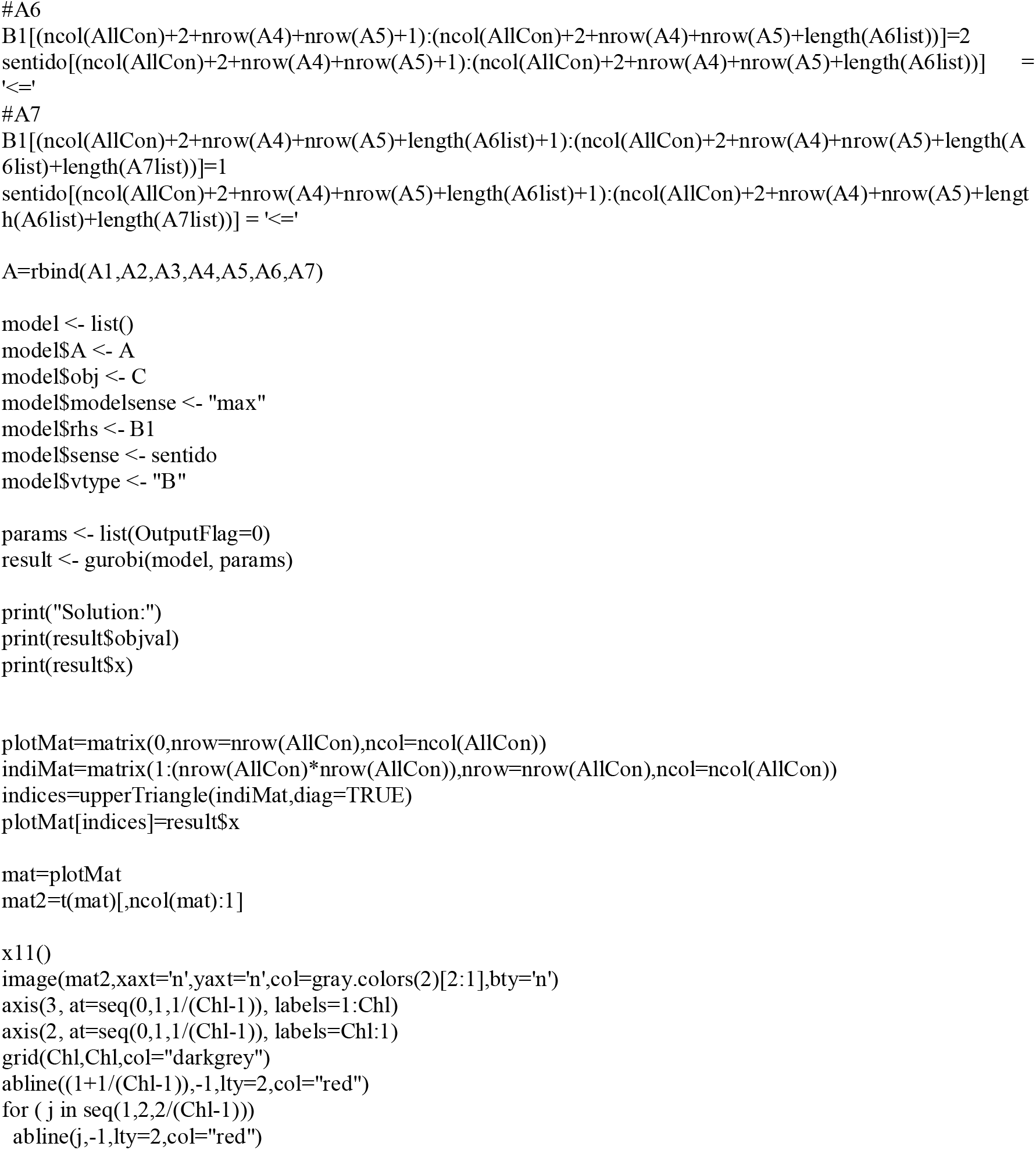

the above code will generate two images of the contact matrix of the globular protein used as example (Fig. S7).

**Fig. S7.**
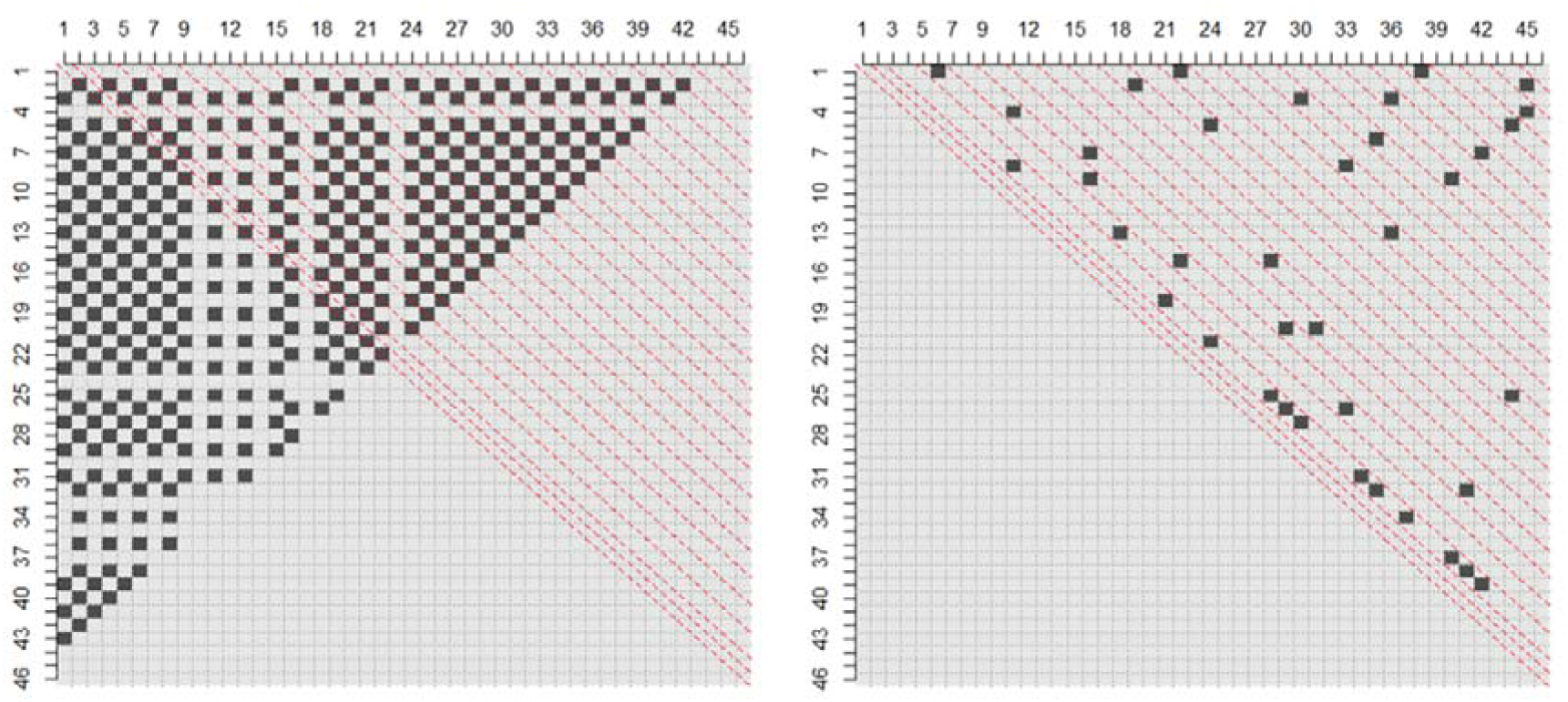
Generation of a protein contact matrix from our simulation. Using the code above and following the instructions above a contact matrix for a 2D representation of a protein is generated: One of the images (left) shows all the possible contacts in the chain and the second image (right) shows the contacts predicted by solving the complex optimization problem.

#### 6.3. Cellular signaling is also a representation of a satisfiability problem (SAT)

Living organisms depend on signaling routes to take care of processes such as survival, apoptosis or tissue repair among many others. This ability of cells to communicate among them is fascinating and constitutes complex networks. To study such complex networks it is common to use semi quantitative computer models that allow to evaluate and characterize different aspects of the net. An example of a cell signaling model is activation and inhibition of platelets in the blood. In the model the arrows indicate a direct activation relation of the node where the arrow starts over the node where the arrow ends. The inhibitory relations are presented as links ending on a perpendicular line and the hierarchy is the same for activation, it means the node where the perpendicular line is, is inhibited by the other node.

**Figure S8:**
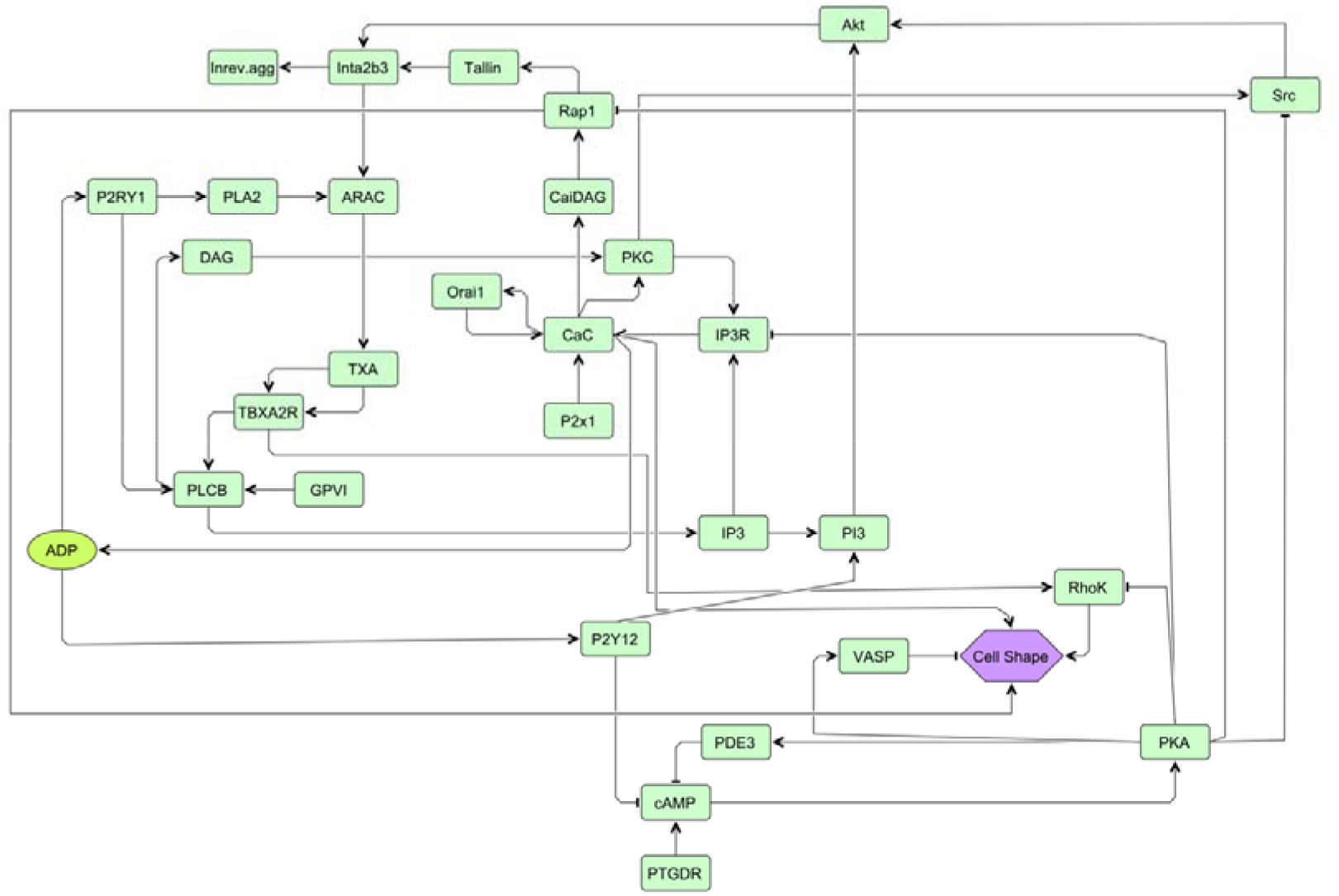
Cell signaling model of Platelet activation. Green boxes represent proteins, their input from other proteins by interactions is shown as activating (arrows) or inhibitory (blunted arrows). Resulting cellular functions are shown as violet hexagons (example: change of cell shape as the first step in platelet activation).

This problem can be stated as a satisfiability problem (SAT), one of the most famous **NP**-complete problems. The Satisfiability problem consists in determining if there exists a combination of variables so a determined Boolean expression is true.

To see how the cell signaling model can be stated as SAT, take for example the cAMP (cyclic adenosine monophosphate) component in the net (Fig. 3). Going step-by-step through the network nodes shown in Fig. 3 (biochemical receptors PTGDR, P2Y12 and molecules activating the cAMP pathway such as phosphodiesterase 3, PDE3 and protein kinase A, PKA as well as calcium in various cellular compartments, CaC, CaE, CaER; details in [5] and considering their logical connectivity we derive then for the nodes a number of expressions such as:

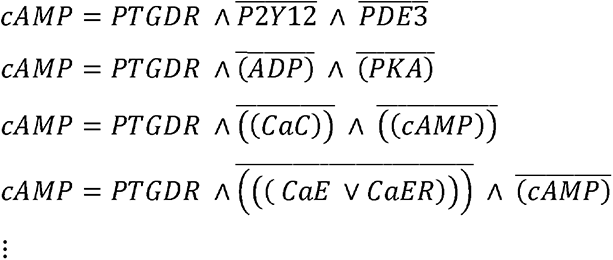

As mentioned before SAT is **NP**-complete although Nature somehow manages to activate the components that need to be activated and inhibits the components that need to be inhibited at the right moments. A famous approximation algorithm for this problem is the Johnson algorithm which basically probes several combinations of variables until finding one that satisfies all parts of the Boolean expression or at least most of them. The search is done in an intelligent manner either by looking to disrupt the less possible of the already satisfied parts or affecting as many as possible among the unsatisfied ones. There is also an approximation algorithm presented by [24] which is a 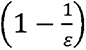-approximation to the optimum.

This algorithm involves a first step of relaxing the SAT problem as a linear programming problem, this relaxation consist in allowing the variables to be relative active, it means the Boolean variables are no longer just zero or one but range in the zero to one interval. Exactly this way occurs in nature where the components of the signaling net are not just fully active or inactive but up to some grade active or inactive. The second step of the algorithm consists in a random process of separation to confirm the membership of components to the active or inactive state, this also happens in nature where everything is a matter of balance and there is a non-rigid threshold which splits between active and inactive components.

The same logical process occurs in metabolic routes in living organism, such as digestion, glycolysis or citric acid cycle among many others.

##### Solution space: going from NP to P

Going from **NP** to **P** can actually mean solving a less limited solution space: when relaxing **NP**-complete problems as shown before what is actually happening is that the feasible solution space is being extended, and not only extended proportional to some constant or even exponentially extended, it is extended to infinite. So if in the worst case scenario of an **NP**-complete problem you have to do an exponential search over all feasible solutions in the worst case scenario of the SDP (Semi-definite Programming) version you would have to do an infinite search through the infinite number of feasible solutions.

For example take the SDP relaxation, it creates a conic feasible region that converts the space from as exponential number of feasible solutions to an infinite number of feasible solutions. The **NP**-complete form of the problem has a finite number of feasible solutions present in the intersections of the restriction planes as presented in **Figure S9**, while the SDP relaxation (the **P** problem) has an infinite number of feasible solutions due to the fact that the cone segment of the feasible region is conformational restricted by an infinite number of restrictive planes so every point in it is a feasible solution.

**Figure S9:**
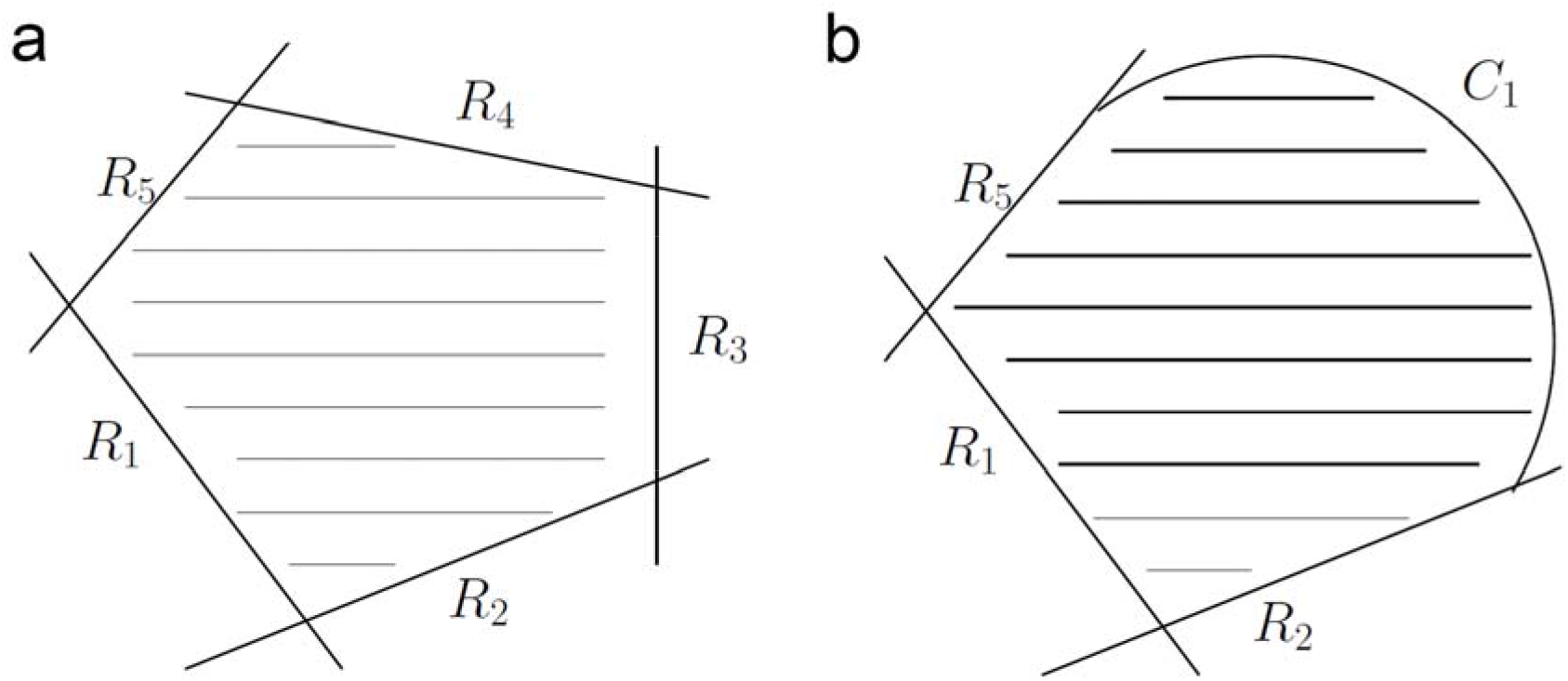
Comparison of feasible regions in NP and P solution spaces. Shown are feasible regions in a) **NP**-complete problem and b) **P** problem form as the SDP (semi-definite programming) relaxation of the **NP**-complete problem. The curved boundary illustrates that the linear boundaries of the solution space are replaced by an infinite number of solutions which can nevertheless be efficiently searched in **P**.

Of course nobody would agree on a claim that **P** might be more complex than **NP** but that is only because we know efficient algorithms to solve problems in **P**. Think for example in the problem of primality (determining whether a number is prime or not) it was not until 2002 that it was proven in **P**, so before that, because of the lack of an efficient algorithm to solve it, it was considered much harder than it is considered today. This might happen also to other problems in the class **NP**, of course the big revolution would be if it ever happens to a problem in **NP**-complete.

We have clearly stated that nature does not solve **NP** problems instead it solves the relaxed version of them. But, what consequences does this relaxation involve? We have seen that the relaxation is always a loose of specificity, it means that things become ambiguous. This ambiguity implies redundancy in every living process, which means that nature, although highly efficient, thanks to evolution and its 3.7 billion years of research and development, is not optimal and not perfect. It can be argued that much of the redundancy found in living processes is intended for security reason and that is true but some of it is also due to the non-optimality of nature. This conjecture can be seen everywhere in nature, from the examples mentioned above to synapsis in neurons which although amazingly efficient is not optimal.

We think nature does not solve **NP**-complete problems in molecular and cellular biology but it certainly does a superb job finding the approximate solutions so we believe that nature has a lot to teach to computer scientists and mathematicians in order to explore new heuristics and new approximations for algorithms development.

The following two Tables illustrate this for decision processes in megakaryocyte maturation and platelet maturation (**Table S6**) and for the protein folding problem (**Table S7**).

#### 6.4. Megakaryocyte and platelet decision processes

We give a lot of data and validation experiments in the following Table S6

**Table S6.** Megakaryocyte platelet formation and platelet signaling as biological decision processes.

##### 6.4.1. Overview: P-problem and no NP-problems involved

1. The robustness in the platelet signaling applies only to the different signaling modules, these are failure tolerant and redundant and robust, only if central hubs are affected, their function stops. Each module is heuristic, has only few nodes and this ensures that there is fast decision in polynomial time (and never anything coming close to an NP problem, in particular there is no unforeseen decision delay)”.
2. Similarly, platelet formation is rather robust (different bystander mutations or KO do not affect at all platelet formation), but if hub is hitting the correct module, then there is thrombocytopenia, but interestingly, the process itself in its speed is NOT affected”

##### 6.4.2. Platelet signaling

Part I: Multiple protein knockouts do not affect the central signaling process, it stays robust and fast. Concrete data that there is a central signaling backbone for efficient signaling transmission and that unless the central backbone is hit, otherwise processing is not harmed, both in mouse and man highly similar and well conserved signaling cascade [12].

Part II: Platelet decision heuristics involves in particular the Src kinase and a bistability switch there to ensure robust, fast and direct decisions. This was confirmed by a combination of light-scattering based thrombocyte aggregation data, western blot and calcium measurements [25].

Evidence 2: Dynamic control centrality describes relaying functions as observed in signalling cascades of the src kinase [26]. The activatory cascade is well conserved and robust, input from neighbor proteins modulates the signal, while the cytoskeleton executes then the platelet activation as a subnetwork [12].

##### 6.4.3. Megakaryocytes and proplatelet formation

Part I: Safe and efficient execution as well as functional independent module s of proplatelet formation, filipodia etc. Modules are here:

a. **cytoplasmic maturation:** STOP (RhoA) and GO (cdc42) function and of course, if you complete smash both parts, thrombocytopenia again. RhoA is also an important target in Myelofibrosis (Fasodil), nice to mention; so this is the affected module, there is no 20-45% proplatelet formation. Upstream of RhoA we have receptors, thrombopoetin and GPIb
b. **Endomitosis** (this is NOT affected, neither by cdc42 KO or RhoA KO): In this module we should mention JAK/STAT pathway, TPO stimulation, normal GATA1 and RunX levels even in the two KO of RhoA and cdc42.
c. **Cytosceletal module**, starting with alpha2-beta1 receptor, tubulin and kindlin, but best we differentiate here also the three steps of platelet activation, intermediate step (alpha2-beta1 receptor) and then more thrombus / 3^rd^ wave.

Part II: Cancer risk rate but parallel to cellular divisions; Evidence: [3, 4].

##### 6.4.4. Further experimental data on the decision modules in platelets and megakaryocytes

The supplementary **Table S3** provides and compares further data for this systems-biological analysis. In particular, specific differences in protein and mRNA expression are listed and compared for strong decision nodes (high centrality values) and unsure decision nodes as well as network connectivity parameters compared. These differences can be rationalized: They optimally support the respective modules in efficient decision processes. Of particular interest is the connectivity for the different types of decision nodes (**Table S3E**) as well as how specific node-specific expression level differences on protein and mRNA level allow that each platelet-specific and also any megakaryocyte-specific module is there in appropriate strength to allow the cell-type specific decision processes and function. In particular, we see that ITGB1 and ILK are clearly undecided nodes and have by far the strongest variance in the female mouse samples. The variance is really high, and all other nodes including the sure nodes have far lower variance. ITGB1 and ILK influence the cytoskeleton and inflammation and, in consequence, lead to the observed myelofibrosis as chronic inflammatory process and output. Hence, this is a good example of the effect of undecided nodes regarding chronic inflammatory signals. Note that on a molecular level, these pathways include cellular decision processes such as cellular signaling including cytoskeletal interactions, cell cycle and apoptosis. The observed chronic inflammation, in this case myelofibrosis, is only a higher level (cellular and tissue processes) of these basic cellular pathway activities.

##### 6.4.5. Further simulation data on the decision modules in platelets and megakaryocytes

**Table S3E** lists the stable system states for the platelet and the HSC network. The nodes are ranked according to their variance of the activation level. The activation level ranges between 0 and 1. The activation level were separated in 10 states between 0 and 1 with a step width of 0.1. It was counted how many different states a node adopts. The nodes that show a high variability of states between 0.1 and 0.9 are called unsure undecided nodes. Nodes that show a high variability but switch between values >0.9 and <0.1 are determined as clearly undecided nodes. The cutoff of variance for undecided nodes was 10^-4^. The Table S3E headers are given here to summarize the overall results from the analysis and the pathway observations made:

- **Sheet I (platelet to megakaryocytes; PL2MK Logfc Female vs Male):** The effect of the KO in G6b on female mice in megakaryocytes in comparison to male mice. Here, SDC4, RBBP8, IKBKB, VAV2, FOXO1, SGK1 are increased by KO in female compared to male and are considered as undecided nodes that change their activity frequently between 0 and 1. Nodes with undecided unsure states are SRF, E2F3, ITGB1, LAT, ILK, RB1, IKBKB and FOXO1. CSK, ITGB1, ILK, FOXO1, IKBKB can be indicated as direct neighbor of the CC in Fig. 2. ITGB1, ILK, IKBKB and FOXO1 are unsure deciders. This shows again that also membrane signaling KOs influence in megakaryocytes platelet generation if the correct signaling modules are hit. The subsequent chronic inflammation leads to myelofibrosis.
- **Sheet II (PL2MK Exprlevel and Variance):** We tried to follow up clearly undecided nodes and unsure undecided nodes in a RNAseq comparison of female and male MK cells and a G6b KO. When analyzing the female vs. male differences in the KO regulation we could outline only slight differences of the mean expression level of unsure nodes and clearly decided nodes. However, when analyzing the variance of expression of unsure nodes compared to clearly decided nodes we could determine a clear difference of variance of the expression level (7,8 vs 0,3). Unsure nodes in general - nodes that are not fully activated nor fully downregulated - show a high variance of the expression data (37,5) compared to the background (9,1) of other proteins. Here, we highlight ITGB1 and ILK that show high variance in the experiment and the simulation, as well as are differential regulated between female and male in the G6b KO study (see also Fig. 2 as 1st neighbor of the central cascade).
- **Sheet III (PL GoEnrichOfUnsureNodes):** The unsure undecided nodes (about 20 in the network; never full active or shut down; high variance in activity between system states, see M&M) regulate specific platelet network modules. The GO (gene ontology) enrichment of the unsure nodes gives an overview of the pathways that are altered by alternative expression of unsure nodes. In the table we recognize that macromolecular metabolic processes in the platelet are influenced by unsure nodes (turnover of RNA in the platelet). mRNA from the megakaryocyte is still present where unsure nodes regulate transcription (see also HSC analysis further columns on the right). The molecular function indicates kinase associations and regulation of nucleic acids. Further less prominent pathways concern related processes in the plasma membrane and cell surface, as well as differentiation and apoptosis. If these processes become unstable they can lead to chronic inflammation. The findings of the *in silico* determined unsure nodes matches with the G6B KO observations: ILK (integrin-linked protein kinase) and ITGB1 (integrin beta 1) are differential regulated in the G6B KO and are indicated as unsure undecided hub nodes in silico, as well as are a close neighbor of the CC it appears, that the KO causes unsure nodes to change their activation level and subsequent sequencing alterations change the cellular state. In an eagle’s perspective the unsure nodes are the effector proteins that are controlled by the central proteins (Fig. 2C) that integrate the cell network information that receptors sensed from the environment. The effector proteins orchestrate the cells response to the environment by changing the expression pattern. Therefore, environmental changes, KO’s and defects are evident of the unsure nodes level with the unsure, flexible, fast, adaptive change in activation and the evidence is also associated with drastic cell state changes, or severe pathologies.
- **Sheet IV (29PI states interpretation):** The platelet has 29 different states in silico. 3 in silico states which are strongly different from each other are exemplified by their predicted network function (KEGG) and the activation. The dominating pathways are here the MAPK pathway that is upregulated in state1, whereas downregulated in state 13 and 29. State 29 is represented by the FoxO signaling activation and state 13 the T cell receptor activation. State 13 seems to activate cancer related proteins. In state 1 the cell cycle is downregulated. Overall, in the platelet quite similar pathways are involved as in the analysis of the HSCs, as again cell cycle and cancer pathways are influenced by the unsure undecided nodes. However, in the mature platelet the pathways for signaling become dominant and are upregulated (MAPK signaling, FoxO signaling, T-cell receptor signaling; KEGG categories are not platelet-specific and hence somewhat crude; note that dominating signal transduction pathways such as src kinase mediated platelet activation are controlled by clearly decided decision nodes. This list concerns only undecided unsure nodes). The unsure undecided nodes influence in the platelet also decisions on platelet signaling in thrombosis and hemostasis implying a risk for cardiovascular pathologies by platelets).
- **Sheet V (HSC GoEnrichOfUnsureNodes):** The unsure undecided nodes (def. see M&M) regulate specific network modules. The GO (gene ontology) enrichment of the unsure nodes gives an overview of the pathways that are altered by the expression of unsure nodes. In the table we find that undecided unsure nodes regulate intracellular signal processes with relation to cardiovascular development. The unsure proteins are located in the cytosol and nucleus and many of them are part of transcription factor complexes. That the unsure nodes of HSC cells - similar to PL unsure nodes - are related to regulation of expression, becomes also clear when analyzing the molecular function: unsure proteins are annotated with protein kinase binding, regulatory regions DNA binding and RNA polymerase binding. Hence, unsure nodes might transmit the decision of the cell network to the DNA expression and thus form a feedback (changing cell state) of the cell on the environmental conditions. The net result is a fine-tuned balance between cell division and cell integrity checks including apoptosis. An inherent risk for imbalance by the unsure undecided nodes implies a cancer risk for HSC (usually leucemias and other blood cancers), or reduced production (myelofibrosis, as also studied in the megakaryocytes by the G6b knockout).
- **Sheet VI (25 HSC states interpretation):** The HSC has 24 different states *in silico*. 3 prominent in silico states are exemplified by their predicted network function (KEGG) and the activation. The dominating pathways are here the MAPK pathway that is upregulated in state 3, 17 and 24 and the p53 pathway, that is downregulated in all investigated states. The pathway analysis show that the states in the HSC are similar in their main pathways, but differ in the weighting of the pathway composition. The inherent error-proneness of the cell division is apparent by several cancer pathways which can be upregulated (state 3, pathways in cancer, endometrial cancer), or downregulated (state 17, state 24). Evidently unsure undecided nodes mediate here a fragile balance between states.

For the platelet the active signal transmission is central, as well as the quiescent state with no activation. Nevertheless, a high number of system states is there, a total of 29 states. To investigate this in more detail (see **Table S3E**), for each state the undecided proteins were divided in two groups: low activity and high activity. Thereby, each state is getting a fingerprint activation pattern of undecided nodes. The pathway each state was analyzed according to its activation pattern. The activation cascade responds to outside signals and uses the decision nodes for turning the system into the activated state. We see here that a fine-tuned activation level is possible (∼70 proteins are high in variance in the simulation, but the others are stable in actvation).

System states are dependent of the number of network modules which is in turn dependent on the network size. Here we have comparable number of interactions (platelet network has 866 interactions and the HSC network has 864 interactions) and thus a comparable number of system states 29, respectively 24 in HSC. the

For comparison, the HSC network is a typical differentiation network, hence plasticity is here necessary, so that than megarkaryozyte, proplatelets and platelets can form. Hence, the good number of system states (24) is reflected here by committed pathways for differentiation. 3 system states stress the activatory cascade already in the megakaryocyte. The commitment to megakaryocyte and platelet formation operates by expression of the suitable modules above and their input to the megakaryocyte system state.

Further references and experimental data on key decision nodes analyzed in the results figures:

###### Central nodes in the platelet network (Fig. 2 in results)

Nodes with high dynamic centrality are known to be key players in experimental literature, such as SRC [5], AKT1 [27, 28], cytosolic calcium (CAC [29]) and ARAC [30, 31].

Finally, cancer risk rate is predicted also for these cell-types to be parallel to cellular divisions (as observed in [3, 4]).

#### 6.5 Protein Folding solutions analysed

Here we give detailed validation data and experiments in Table S7.

**Table S7.**
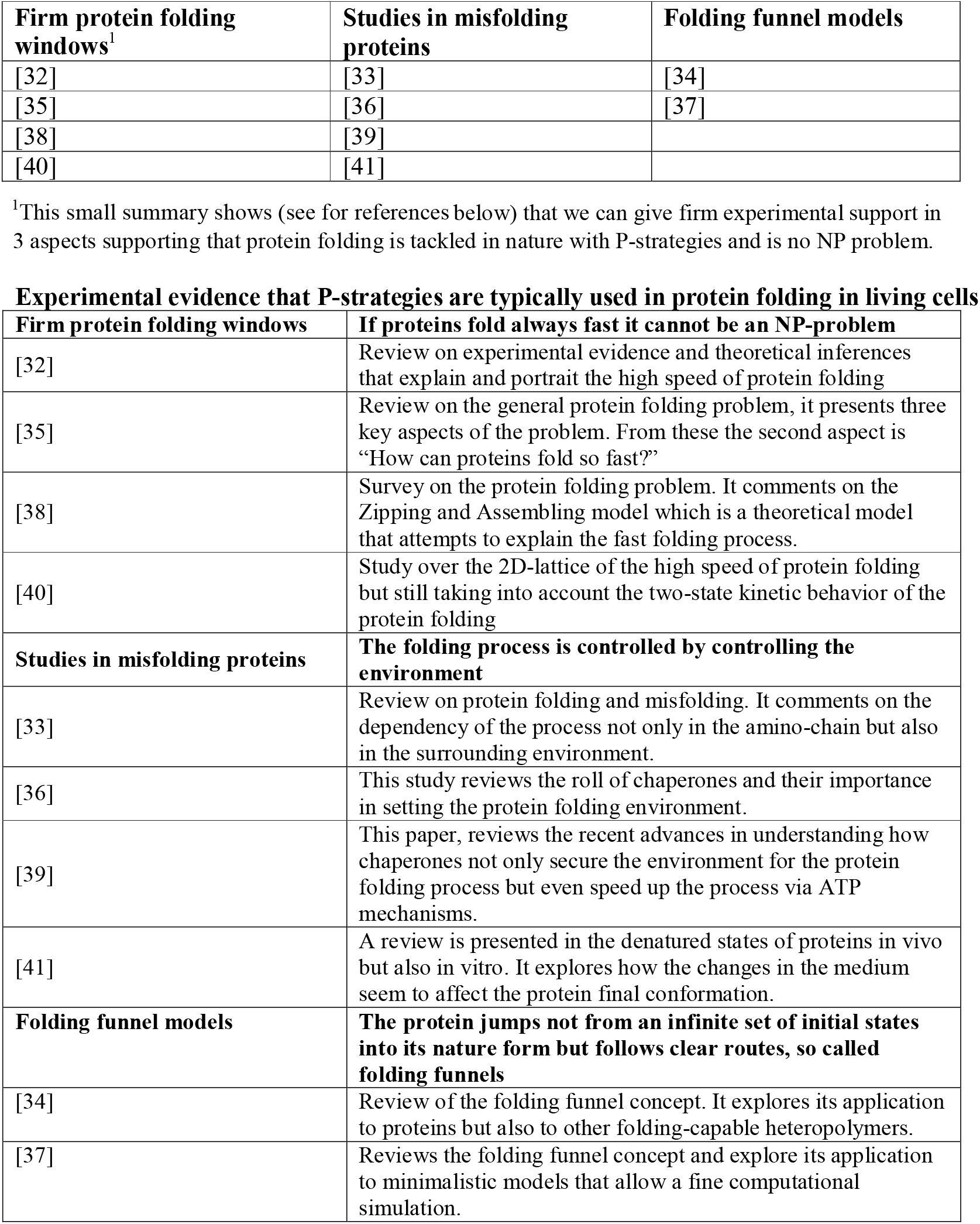
Nature uses only P-strategies in Protein folding.

### 7. On the relation of P and NP problems

As stated by Stephen Cook, “The P vs NP problem is to determine whether every language accepted by some nondeterministic Turing machine in polynomial time is also accepted by some deterministic Turing machine in polynomial time”. This is, whether is it possible to use a computer to efficiently (polynomial time) solve problems which can be efficiently solved by an omniscient oracle. Hence, in principle, what is being asked there is whether there exist a set of problems which are essentially harder than others and which cannot be solved efficiently using a deterministic machine.

The general consensus nowadays is that P is not equal NP, but so far this has not been proved. The same Stephen Cook in a paper called “The importance of the P vs NP question” debates over three possible outcomes for this question.

1. Somebody will prove that P is different to NP. In this case things do not change much, we still not have efficient solutions for NP complete problems and will never have, more effort would be spent in sub-optimal efficient solutions.
2. P is proved to be equal to NP in a non-constructive way. This means that they are known to be equal but no mapping is known between them or the known mapping is polynomial with a really high grade. In this case things also do not change much. NP complete problems remain intractable for practical purposes but anyway the search for polynomial algorithms for those problems is strongly fostered.
3. Last possible outcome, an efficient mapping is found from P to NP. All NP complete problems are now solved in polynomial time. This is a major change for society, all mathematical proves can now be done by a machine. Artificial intelligence becomes a reality. And finally, all digital security systems need to be changed, nowadays cryptography becomes useless.

As can be seen in the poll from William Gasarch from 2002, where 61 experts replied that P=NP will be proved someday against only 9 that think that the opposite will be proved, the general believe over time[42] is that there is a certain kind of problems which cannot be solved efficiently. Here we sum ourselves to the group of researchers that believe in the “P not-equal NP” outcome and we bring with us all living organisms together with their millions of years in research and development through evolution. We claim that nature when facing NP problems opts for a pragmatic approach and settles for a sub-optimal solution of the problem. Through years of refinement and with the luxury of no deadline and no limit of test subjects Nature has come to the point of finding remarkable close to the optimum solutions to many NP problems found in living organisms, but nevertheless these solutions remain suboptimal. We see how for example in protein folding the native conformation of the proteins is achieved through a delicate balance in all conditions where the folding takes place, even to the point of using “folding cameras” (chaperon proteins) where the protein is free from obstructions. This ensures that the folding process starts always from the same point so it ends up always in the same local minimum, but there is absolutely no guarantee of finding the absolute minimum in the funnel landscape.

This pragmatic approach from living organisms is somehow similar to the non-deterministic Turing machine. Living organisms are often impelled to take decisions without full knowledge of the problem or without enough time to solve it. This leads to suboptimal solutions but in most cases good enough for the main purpose of life, survival. We see examples of this in the platelet and the stems-cell networks we present.

However, regarding physics, limitations of biology sometimes may play no role, for instance in quantum physics, hence here more demanding computations are possible. We are aware that even a quantum computer may not allow to solve all NP problems: it efficiently solves factoring and discrete logarithm calculations, however, graph isomorphisms and NP complete problems may still pose a challenge [23]; this is illustrated in Fig. 3e in the paper. However, and in contrast, natural physical processes really emulate all combinatorial events and paths in parallel, for instance the Feynman path integral traces all energy levels and trajectories. Only by this massive inbuilt quantum parallelism we are able to find the observed trajectory and so here the complete problem space is covered for quantum processes.

#### Our basic assumption: NP problems never can be mapped to P problems

Our suggestion is to check the following arguments in mathematical detail in search for such a firm proof:

Cantoŕs Diagonal proof shows that no mapping of R to Q or N is possible. However, this is an uncountable infinite set, so really huge, the set of R, so no wonder, that you easily show no mapping there to the smaller set of countable numbers.

However, something related to the Diagonal proof could may be apply to the NP to P problem:

The challenge is that an individual solution to an NP problem is easily checked for its result (e.g. in the TSP), however, you never know whether this is already the best solution.

Hence, this means, you have **no ordered dictionary of solutions.**

We think this is already clear, in a P problem you have sorted solutions and so you easily know, well, this is still far away from the best solution.

but in NP there is no such sorting there, so you never know whether it is the best solution or not (may be this statement for some self-evident).

However, our proof conjecture would then be:

If it is known for one set of problems there is no sorting of the solutions possible (problem is known to be NP or at least part of the NP complete problem class; e.g. ranked according to energy/distance, more accurately by looking at the solution you DO NOT KNOW that this is the best solution or very far from the minimum) and there are classical P problems, where there is a sorted dictionary IS possible.

Then it should be possible (e.g by an indirect proof or something like this) that you can proof no mapping of the NOT sortable solution space to the problem class of SORTED solutions is possible.

In conclusion, the sorted dictionary of solutions property alone not only distinguishes both problem classes but we can derive from this property maybe a proof that no mapping between **NP** and **P** problems is possible.

## References

1. Cook S. The P vs NP problem. Commun ACM. 2009. doi:10.1145/1562164.1562186

2. Gasarch W. Guest Column: The Third P=?NP Poll. ACM SIGACT News. 2019;50: 38–59. doi:10.1145/3319627.3319636

3. Raatikainen P. Gödel’s Incompleteness Theorems. Fall 2018. In: Zalta EN, editor, The Stanford Encyclopedia of Philosophy. Fall 2018. Metaphysics Research Lab, Stanford University; 2018.

4. Tomasetti C, Durrett R, Kimmel M, Lambert A, Parmigiani G, Zauber A, et al. Role of stem-cell divisions in cancer risk. Nature. 2017;548: E13–E14. doi:10.1038/nature23302

5. Jumper J, Evans R, Pritzel A, Green T, Figurnov M, Ronneberger O, et al. Highly accurate protein structure prediction with AlphaFold. Nat 2021. 2021; 1–11. doi:10.1038/s41586-021-03819-2

6. Goemans MX, Williamson DP. Approximation algorithms for Max-3-Cut and other problems via complex semidefinite programming. J Comput Syst Sci. 2004;68: 442– 470. doi:10.1016/j.jcss.2003.07.012

7. Vaquer-Alicea J, Diamond MI. Propagation of Protein Aggregation in Neurodegenerative Diseases. Annu Rev Biochem. 2019. doi:10.1146/annurev-biochem-061516-045049

8. Sergiev P V., Dontsova OA, Berezkin G V. Theories of aging: An ever-evolving field. Acta Naturae. Russian Federation Agency for Science and Innovation; 2015. pp. 9–18. doi:10.32607/20758251-2015-7-1-9-18

9. Troelstra AS. Aspects of Constructive Mathematics. Stud Log Found Math. 1977;90: 973–1052. doi:10.1016/S0049-237X(08)71127-3

10. Hofstadter DR. Go□del, Escher, Bach□: an eternal goldern braid. Basic Books; 1979.

11. Chaitin G. The limits of reason. Scientific American. Scientific American Inc.; 2006. pp. 74–81. doi:10.1038/scientificamerican0306-74

12. Tomasetti C, Li L, Vogelstein B. Stem cell divisions, somatic mutations, cancer etiology, and cancer prevention. Science (80- ). 2017;355: 1330–1334. doi:10.1126/science.aaf9011

13. Berger B, Leighton T. Protein folding in the hydrophobic-hydrophilic (HP) model is NP-complete. Proceedings of the Annual International Conference on Computational Molecular Biology, RECOMB. New York, New York, USA: ACM; 1998. pp. 30–39. doi:10.1145/279069.279080

14. Dill KA, MacCallum JL. The protein-folding problem, 50 years on. Science (80- ). 2012;338: 1042–1046. doi:10.1126/science.1219021

15. Goemans MX, Williamson DP. Improved Approximation Algorithms for Maximum Cut and Satisfiability Problems Using Semidefinite Programming. J ACM. 1995;42: 1115–1145. doi:10.1145/227683.227684

16. Chen J, Schafer NP, Wolynes PG, Clementi C. Localizing Frustration in Proteins Using All-Atom Energy Functions. J Phys Chem B. 2019;123: 4497–4504. doi:10.1021/acs.jpcb.9b01545

17. Truong HH, Kim BL, Schafer NP, Wolynes PG. Predictive energy landscapes for folding membrane protein assemblies. J Chem Phys. 2015;143: 243101. doi:10.1063/1.4929598

18. Bar-Lavan Y, Shemesh N, Ben-Zvi A. Chaperone families and interactions in metazoa. Essays Biochem. 2016;60: 237–253. doi:10.1042/EBC20160004

19. Partridge L, Barton NH. Optimally, mutation and the evolution of ageing. Nature. 1993;362: 305–311. doi:10.1038/362305a0

20. Ocampo A, Reddy P, Martinez-Redondo P, Platero-Luengo A, Hatanaka F, Hishida T, et al. In Vivo Amelioration of Age-Associated Hallmarks by Partial Reprogramming. Cell. 2016;167: 1719–1733.e12. doi:10.1016/j.cell.2016.11.052

21. Penrose R. Shadows of the Mind: A Search for the Missing Science of Consciousness. Oxford University Press; 1994.

22. Boyanova D, Nilla S, Birschmann I, Dandekar T, Dittrich M. PlateletWeb: A systems biologic analysis of signaling networks in human platelets. Blood. 2012;119. doi:10.1182/blood-2011-10-387308

23. Balkenhol J, Kaltdorf K V., Mammadova-Bach E, Braun A, Nieswandt B, Dittrich M, et al. Comparison of the central human and mouse platelet signaling cascade by systems biological analysis. BMC Genomics. 2020;21. doi:10.1186/s12864-020-07215-4

24. Remmele CW, Luther CH, Balkenhol J, Dandekar T, Müller T, Dittrich MT. Integrated inference and evaluation of host-fungi interaction networks. Front Microbiol. 2015;6: 764. doi:10.3389/fmicb.2015.00764

25. Ramström S. Arachidonic acid causes lysis of blood cells and ADP-dependent platelet activation responses in platelet function tests. Platelets. 2019;30: 1001–1007. doi:10.1080/09537104.2018.1557614

26. Sangkuhl K, Shuldiner AR, Klein TE, Altman RB. Platelet aggregation pathway. Pharmacogenet Genomics. 2011;21: 516–521. doi:10.1097/FPC.0b013e3283406323

27. Woulfe DS. Akt signaling in platelets and thrombosis. Expert Review of Hematology. 2010. pp. 81–91. doi:10.1586/ehm.09.75

28. Yin H, Stojanovic A, Hay N, Du X. The role of Akt in the signaling pathway of the glycoprotein Ib-IX-induced platelet activation. Blood. 2008;111: 658–665. doi:10.1182/blood-2007-04-085514

29. Mischnik M, Boyanova D, Hubertus K, Geiger J, Philippi N, Dittrich M, et al. A Boolean view separates platelet activatory and inhibitory signalling as verified by phosphorylation monitoring including threshold behaviour and integrin modulation. Mol Biosyst. 2013;9: 1326–1339. doi:10.1039/c3mb25597b

30. Rowley JW, Oler AJ, Tolley ND, Hunter BN, Low EN, Nix DA, et al. Genome-wide RNA-seq analysis of human and mouse platelet transcriptomes. Blood. 2011;118: e101. doi:10.1182/blood-2011-03-339705

31. The UniProt Consortium, Bateman A, Martin M-J, Orchard S, Magrane M, Agivetova R, et al. UniProt: the universal protein knowledgebase in 2021. Nucleic Acids Res. 2021;49: D480–D489. doi:10.1093/NAR/GKAA1100

32. Shannon P, Markiel A, Ozier O, Baliga NS, Wang JT, Ramage D, et al. Cytoscape: A software Environment for integrated models of biomolecular interaction networks. Genome Res. 2003. doi:10.1101/gr.1239303

33. Mi H, Huang X, Muruganujan A, Tang H, Mills C, Kang D, et al. PANTHER version 11: Expanded annotation data from Gene Ontology and Reactome pathways, and data analysis tool enhancements. Nucleic Acids Res. 2017. doi:10.1093/nar/gkw1138

34. Karl S, Dandekar T. Convergence behaviour and Control in Non-Linear Biological Networks. Sci Rep. 2015;5: 1–11. doi:10.1038/srep09746

35. Karl S, Dandekar T. Jimena: Efficient computing and system state identification for genetic regulatory networks. BMC Bioinformatics. 2013;14: 306. doi:10.1186/1471-2105-14-306

36. Mischnik M, Gambaryan S, Subramanian H, Geiger J, Schütz C, Timmer J, et al. A comparative analysis of the bistability switch for platelet aggregation by logic ODE based dynamical modeling. Mol Biosyst. 2014;10: 2082–2089. doi:10.1039/c4mb00170b

37. Molania R, Gagnon-Bartsch JA, Dobrovic A, Speed TP. A new normalization for Nanostring nCounter gene expression data. Nucleic Acids Res. 2019;47: 6073–6083. doi:10.1093/nar/gkz433

38. Chen Y, McCarthy D, Ritchie M, Robinson M, Smyth G, Hall E. edgeR: differential analysis of sequence read count data User’s Guide. R Packag. 2020; 1–121.

39. Hochberg Y, Benjamini Y. More powerful procedures for multiple significance testing. Stat Med. 1990;9: 811–818. doi:10.1002/sim.4780090710

40. Hetz C, Saxena S. ER stress and the unfolded protein response in neurodegeneration. Nature Reviews Neurology. Nature Publishing Group; 2017. pp. 477–491. doi:10.1038/nrneurol.2017.99

41. Aaronson S. The limits of quantum computers. Lecture Notes in Computer Science (including subseries Lecture Notes in Artificial Intelligence and Lecture Notes in Bioinformatics). Springer, Berlin, Heidelberg; 2007. p. 4. doi:10.1007/978-3-540-74510-5_2

42. Borwein J, Bailey D. Mathematics by experiment, 2nd edition: Plausible reasoning in the 21st century. Mathematics by Experiment, 2nd Edition: Plausible Reasoning in the 21st Century. 2008.

43. Felder K. Kenny’s overview of Hofstadter’s Explanation of Godel’s theorem, the website. In: http://www.felderbooks.com/papers/godel. 1996.

## References

1. Cook SA. The complexity of theorem-proving procedures. Proceedings of the Annual ACM Symposium on Theory of Computing. New York, New York, USA: Association for Computing Machinery; 1971. pp. 151–158. doi:10.1145/800157.805047

2. Levin L. Universal sequential search problems. Probl Peredachi Informatsii. 1973.

3. Tomasetti C, Li L, Vogelstein B. Stem cell divisions, somatic mutations, cancer etiology, and cancer prevention. Science (80- ). 2017;355: 1330–1334. doi:10.1126/science.aaf9011

5. Mischnik M, Boyanova D, Hubertus K, Geiger J, Philippi N, Dittrich M, et al. A Boolean view separates platelet activatory and inhibitory signalling as verified by phosphorylation monitoring including threshold behaviour and integrin modulation. Mol Biosyst. 2013;9: 1326–1339. doi:10.1039/c3mb25597b

6. Hofstadter DR. Go□del, Escher, Bach□: an eternal goldern braid. Basic Books; 1979.

7. Chaitin G. The limits of reason. Scientific American. Scientific American Inc.; 2006. pp. 74–81. doi:10.1038/scientificamerican0306-74

8. Vastrik I, D’Eustachio P, Schmidt E, Joshi-Tope G, Gopinath G, Croft D, et al. Reactome: A knowledge base of biologic pathways and processes. Genome Biol. 2007. doi:10.1186/gb-2007-8-3-r39

9. Mukherji M, Bell R, Supekova L, Wang Y, Orth AP, Batalov S, et al. Genome-wide functional analysis of human cell-cycle regulators. Proc Natl Acad Sci U S A. 2006;103: 14819–14824. doi:10.1073/pnas.0604320103

10. Ubersax JA, Woodbury EL, Quang PN, Paraz M, Blethrow JD, Shah K, et al. Targets of the cyclin-dependent kinase Cdk1. Nature. 2003. doi:10.1038/nature02062

11. Czakai K, Dittrich M, Kaltdorf M, Müller T, Krappmann S, Schedler A, et al. Influence of Platelet-rich Plasma on the immune response of human monocyte-derived dendritic cells and macrophages stimulated with Aspergillus fumigatus. Int J Med Microbiol. 2017;307: 95–107. doi:10.1016/j.ijmm.2016.11.010

12. Balkenhol J, Kaltdorf K V., Mammadova-Bach E, Braun A, Nieswandt B, Dittrich M, et al. Comparison of the central human and mouse platelet signaling cascade by systems biological analysis. BMC Genomics. 2020;21. doi:10.1186/s12864-020-07215-4

13. Breitenbach T, Liang C, Beyersdorf N, Dandekar T. Analyzing pharmacological intervention points: A method to calculate external stimuli to switch between steady states in regulatory networks. Papin JA, editor. PLOS Comput Biol. 2019;15: e1007075. doi:10.1371/journal.pcbi.1007075

14. Yang JH, Wright SN, Hamblin M, McCloskey D, Alcantar MA, Schrübbers L, et al. A White-Box Machine Learning Approach for Revealing Antibiotic Mechanisms of Action. Cell. 2019;177: 1649–1661.e9. doi:10.1016/j.cell.2019.04.016

15. Naseem M, Othman EM, Fathy M, Iqbal J, Howari FM, AlRemeithi FA, et al. Integrated structural and functional analysis of the protective effects of kinetin against oxidative stress in mammalian cellular systems. Sci Rep. 2020;10: 1–10. doi:10.1038/s41598-020-70253-1

16. Srivastava M, Bencurova E, Gupta SK, Weiss E, Löffler J, Dandekar T. Aspergillus fumigatus challenged by human dendritic cells: Metabolic and regulatory pathway responses testify a tight battle. Front Cell Infect Microbiol. 2019;9: 168. doi:10.3389/fcimb.2019.00168

17. Goemans MX, Williamson DP. Approximation algorithms for Max-3-Cut and other problems via complex semidefinite programming. J Comput Syst Sci. 2004;68: 442–470. doi:10.1016/j.jcss.2003.07.012

18. Hetz C, Saxena S. ER stress and the unfolded protein response in neurodegeneration. Nature Reviews Neurology. Nature Publishing Group; 2017. pp. 477–491. doi:10.1038/nrneurol.2017.99

19. Schaarschmidt J, Monastyrskyy B, Kryshtafovych A, Bonvin AMJJ. Assessment of contact predictions in CASP12: Co-evolution and deep learning coming of age. Proteins Struct Funct Bioinforma. 2018;86: 51–66. doi:10.1002/prot.25407

20. Weisstein EW. CRC Concise Encyclopedia of Mathematics. CRC Concise Encyclopedia of Mathematics. 2002. doi:10.1201/9781420035223

21. Baker A. Transcendental Number Theory. Cambridge University Press; 1975. doi:10.1017/CBO9780511565977

22. Penrose R. Shadows of the Mind: A Search for the Missing Science of Consciousness. Oxford University Press; 1994.

23. Aaronson S. The limits of quantum computers. Lecture Notes in Computer Science (including subseries Lecture Notes in Artificial Intelligence and Lecture Notes in Bioinformatics). Springer, Berlin, Heidelberg; 2007. p. 4. doi:10.1007/978-3-540-74510-5_2

24. Goemans MX, Williamson DP. Improved Approximation Algorithms for Maximum Cut and Satisfiability Problems Using Semidefinite Programming. J ACM. 1995;42: 1115–1145. doi:10.1145/227683.227684

25. Mischnik M, Gambaryan S, Subramanian H, Geiger J, Schütz C, Timmer J, et al. A comparative analysis of the bistability switch for platelet aggregation by logic ODE based dynamical modeling. Mol Biosyst. 2014;10: 2082–2089. doi:10.1039/c4mb00170b

26. Karl S, Dandekar T. Convergence behaviour and Control in Non-Linear Biological Networks. Sci Rep. 2015;5: 1–11. doi:10.1038/srep09746

29. Varga-Szabo D, Braun A, Nieswandt B. Calcium signaling in platelets. Journal of Thrombosis and Haemostasis. 2009. doi:10.1111/j.1538-7836.2009.03455.x

30. Ramström S. Arachidonic acid causes lysis of blood cells and ADP-dependent platelet activation responses in platelet function tests. Platelets. 2019;30: 1001–1007. doi:10.1080/09537104.2018.1557614

31. Sangkuhl K, Shuldiner AR, Klein TE, Altman RB. Platelet aggregation pathway. Pharmacogenet Genomics. 2011;21: 516–521. doi:10.1097/FPC.0b013e3283406323

32. Gelman H, Gruebele M. Fast protein folding kinetics. Q Rev Biophys. 2014;47: 95–142. doi:10.1017/S003358351400002X

33. Dobson CM. Protein folding and misfolding. Nature. Nature Publishing Group; 2003. pp. 884–890. doi:10.1038/nature02261

34. Onuchic JN, Luthey-Schulten Z, Wolynes PG. Theory of Protein Folding: The Energy Landscape Perspective. Annu Rev Phys Chem. 1997;48: 545–600. doi:10.1146/annurev.physchem.48.1.545

35. Dill KA, MacCallum JL. The protein-folding problem, 50 years on. Science (80- ). 2012;338: 1042–1046. doi:10.1126/science.1219021

36. Kim YE, Hipp MS, Bracher A, Hayer-Hartl M, Ulrich Hartl F. Molecular chaperone functions in protein folding and proteostasis. Annual Review of Biochemistry. Annu Rev Biochem; 2013. pp. 323–355. doi:10.1146/annurev-biochem-060208-092442

37. Onuchic JN, Wolynes PG. Theory of protein folding. Current Opinion in Structural Biology. Elsevier Ltd; 2004. pp. 70–75. doi:10.1016/j.sbi.2004.01.009

38. Dill KA, Ozkan SB, Shell MS, Weikl TR. The protein folding problem. Annual Review of Biophysics. NIH Public Access; 2008. pp. 289–316. doi:10.1146/annurev.biophys.37.092707.153558

39. Balchin D, Hayer-Hartl M, Hartl FU. Recent advances in understanding catalysis of protein folding by molecular chaperones. FEBS Lett. 2020;594: 2770–2781. doi:10.1002/1873-3468.13844

40. Schonbrun J, Dill KA. Fast protein folding kinetics. Proc Natl Acad Sci U S A. 2003;100: 12678–12682. doi:10.1073/pnas.1735417100

41. Dill KA. Denatured states of proteins. Annual Review of Biochemistry. Annual Reviews Inc.; 1991. pp. 795–825. doi:10.1146/annurev.bi.60.070191.004051

42. Gasarch W. Guest Column: The Third P=?NP Poll. ACM SIGACT News. 2019;50: 38–59. doi:10.1145/3319627.3319636

